# Distentangling the systems contributing to changes in learning during adolescence

**DOI:** 10.1101/622860

**Authors:** Sarah L. Master, Maria K. Eckstein, Neta Gotlieb, Ronald Dahl, Linda Wilbrecht, Anne G.E. Collins

## Abstract

Multiple neurocognitive systems contribute simultaneously to learning. For example, dopamine and basal ganglia (BG) systems are thought to support reinforcement learning (RL) by incrementally updating the value of choices, while the prefrontal cortex (PFC) contributes different computations, such as actively maintaining precise information in working memory (WM). It is commonly thought that WM and PFC show more protracted development than RL and BG systems, yet their contributions are rarely assessed in tandem. Here, we used a simple learning task to test how RL and WM contribute to changes in learning across adolescence. We tested 187 subjects ages 8 to 17 and 53 adults (25-30). Participants learned stimulus-action associations from feedback; the learning load was varied to be within or exceed WM capacity. Participants age 8-12 learned slower than participants age 13-17, and were more sensitive to load. We used computational modeling to estimate subjects’ use of WM and RL processes. Surprisingly, we found more robust changes in RL than WM during development. RL learning rate increased significantly with age across adolescence and WM parameters showed more subtle changes, many of them early in adolescence. These results underscore the importance of changes in RL processes for the developmental science of learning.

**Highlights:**

- Subjects combine reinforcement learning (RL) and working memory (WM) to learn
- Computational modeling shows RL learning rates grew with age during adolescence
- When load was beyond WM capacity, weaker RL compensated less in younger adolescents
- WM parameters showed subtler and more puberty-related changes
- WM reliance, maintenance, and capacity had separable developmental trajectories
- Underscores importance of RL processes in developmental changes in learning

## Introduction

There is increasing evidence that multiple neural systems contribute to human learning ((Hazy, Frank, & O’Reilly, 2007; Myers et al., 2002)), even in simple cognitive paradigms previously modeled with a single learning process ((Bornstein & Daw, 2012; Bornstein & Norman, 2017; Collins & Frank, 2012)). Basal ganglia (BG) dependent reinforcement learning (RL) processes are thought to be supplemented by multiple other systems, including prefrontal executive functions ((Badre, Kayser, & D’Esposito, 2010)) such as working memory (WM; (Collins & Frank, 2012)) and model-based planning ((Daw et al., 2011)), as well as hippocampus-based episodic memory ((Bornstein & Daw, 2012; Bornstein & Norman, 2017; Davidow et al., 2016; Myers et al., 2002; Wimmer et al., 2014)). To understand developmental changes in learning, it is important to carefully capture the contributions of these multiple systems to learning. Previous work has shown differential developmental trajectories for RL, episodic memory, and model-based planning (Crone, et al., 2006; Decker et al., 2016; Selmeczy et al., 2018; Somerville, Jones, & Casey, 2010). Here, we investigate how the relative contributions of RL and WM change during development.

RL is an incremental learning process which updates stored choice values from the discrepancy between obtained and expected reward (the reward prediction error; RPE) in order to maximize future rewards. Sensitivity to rewarding outcomes in the ventral striatum has been shown to peak during mid-adolescence, relative to children and adults ((Galvan et al., 2006; Braams et al., 2015; Gulley et al., 2018; Somerville et al., 2010)). It has been proposed that the BG are ‘mature’ by mid adolescence, but it is also known that structures that provide inputs to the BG continue to show anatomical and functional change ((Braams, van Duijvenvoorde, Peper, & Crone, 2015; Casey et al., 2008; A. Galvan et al., 2006a; Galvan, Delevich, & Wilbrecht, 2019; Van Den Bos, Cohen, Kahnt, & Crone, 2012)).

The prefrontal cortex (PFC) is known to develop late into adolescence and the third decade of life ((Casey et al., 2008; Giedd, Blumenthal, & Jeffries, 1999; Larsen & Luna, 2018)) and is thought to be critical for WM performance ((Curtis & D’Esposito, 2003; Miller & Cohen, 2001)). WM allows for the immediate and accurate storage of information, but representations in WM are thought to decay quickly with time and are subject to interference, as the information that can be held in WM is limited ((D’Esposito & Postle, 2015; Oberauer et al., 2018)). Therefore there may be tradeoffs to using fast capacity-limited WM, versus slower, capacity unlimited RL-based learning systems.

Behavioral testing and computational modeling can be used to disentangle the roles of RL and WM in human learning, and how their use differs between individuals. Using a deterministic reward-learning task called “RLWM” that taxes WM by varying the amount of information to learn in each block, we have previously isolated contributions of WM and RL learning ((Collins, 2018; Collins & Frank, 2012, 2018)). In multiple studies we found that participants mainly used WM for learning when the load was within their capacity, and otherwise compensated with RL. We also found that learning deficits in schizophrenia were a result of weakened WM contributions with intact RL ((Collins & Frank, 2014; Collins et al., 2017). Here we used the same approach to investigate the maturation of WM and RL and their relative contribution to learning across adolescent development (sampling subjects 8-17 and 25-30).

Using behavioral testing and computational modeling, we examined three separate hypotheses of how RL-and WM-based learning develop relative to each other. Our first hypothesis was that both RL and WM systems’ contributions to learning show continued development into later adolescence such that both systems are dynamic throughout the teenage years. Developmental changes in WM are prominent in the literature on development of executive function, including the maintenance ((Geier et al., 2009; McAuley & White, 2011)) and manipulation of information in WM ((Crone et al., 2006; Huizinga, Dolan, & van der Molen, 2006)), as well as the precision of the representations in WM ((Luna, 2009)). There is also a strong literature showing changes in RL learning systems from childhood to adulthood, such as dynamic changes in reward sensitivity in the striatum across adolescence ((Braams et al., 2015; Cohen et al., 2010; Davidow et al., 2016; Somerville et al., 2010)) and changes in learning rate between adolescence and adulthood ((Davidow et al., 2016)).

Our second hypothesis emphasized the relative importance of WM development over that of RL in accounting for changes in learning in adolescence. Dual systems models and other popular models of adolescent development place great weight on the late maturation of the PFC and PFC-dependent executive functions, such as WM or model-based learning ((Casey et al., 2008; Decker et al., 2016; Huizinga et al., 2006; Steinberg, 2005)). Our second hypothesis, therefore predicts that even thought RL may be developing, we should observe stronger effects of age on WM, which is PFC-based, and WM changes should be the primary driver of changes in learning through adolescence.

Finally, we hypothesized that pubertal onset may significantly impact WM processes. There is growing evidence that gonadal hormones affect inhibitory neurotransmission and other variables in the prefrontal cortex of rodents (Delevich, Piekarski, & Wilbrecht, 2019; Delevich, Thomas, & Wilbrecht, 2019; Juraska & Willing, 2017; Piekarski, Boivin, & Wilbrecht, 2017; Piekarski, Johnson, et al., 2017). We therefore predicted that WM parameters would differ in children with different pubertal status or gonadal hormone concentration.

To evaluate these hypotheses, we tested children and adolescents aged 8 to 17 years old and adults aged 25 to 30 years old on the RLWM task (Collins & Frank, 2012). We then fit computational models of behavior to subjects’ performance and assessed how these parameters changed with age, pubertal development and salivary testosterone levels. Using these established methods to disentangle the contributions of RL and WM, we found changes in RL spanning adolescent development, but much weaker changes in WM contributions. WM differences did show relationships with pubertal variables.

Overall, these data support the somewhat surprising conclusion that changes in RL systems are important drivers of change in simple associative learning during adolescence. The results also support further inquiry into the role of pubertal processes in WM function.

## Methods

### Subject testing

All procedures were approved by the Committee for the Protection of Human Subjects at the University of California, Berkeley. After entering the testing room, subjects under 18 years old and their guardians provided their informed assent or permission. All guardians were asked to fill out a demographic form, and guardians of children aged 10 or younger completed an additional pubertal development questionnaire about their child. Subjects were led into a quiet testing room in view of their guardians, where they used a video game controller to complete four computerized tasks. An hour after the start of the experimental session and in between tasks, subjects provided a 1.8 mL saliva sample. At the conclusion of the tasks, subjects were asked to complete a short questionnaire which assessed their pubertal development ((Petersen & Crockett, 1988)) and collected other basic information, like their height and weight. Subjects under 10 years old did not complete the pubertal development questionnaire. Subjects were then compensated with one $25 Amazon gift card for their time.

Participants over 18 provided informed consent and completed all forms themselves. They also answered retroactive questions about puberty, otherwise all testing procedures were identical.

### Demographics

We recruited 191 children and 55 adults to participate in this study. Out of those who reported their race, 60 subjects identified as Asian, 10 African-American, and 6 Native American or Pacific Islander. 28 were of mixed race. The remaining 127 subjects identified as Caucasian. 29 subjects identified as Hispanic. 4 children and 1 adult failed to complete the task, either out of disinterest or because of controller malfunction. All subjects who completed the task performed above chance (33% accuracy), so none were excluded for poor task performance. 187 children (89 female, mean (std) age 12.62 (2.76) years) and 54 adults (28 female, mean (std) age 26.77 (1.49) years) were included in analyses.

### Experimental paradigm

The experiment described in this work was the second of four administered tasks. The preceding task was a 5 to 10 minute deterministic learning task in which subjects learned to select one correct action out of four possible actions. Halfway through, the correct action switched. The results for this and the other tasks will be reported elsewhere.

This experiment is based on the “RLWM” task described in ((Collins, Ciullo, Frank, & Badre, 2017; Collins & Frank, 2012, 2018)), adapted for the developmental population. To make the task more engaging for children, participants were told that aliens had landed on earth and wanted to teach us their alien language. In this alien language, one button on the controller was matched with each image on the screen. Participants completed one block of training and then ten independent learning blocks, for a total duration of less than 25 minutes (mean duration 16.9 minutes, range 14 – 25 minutes).

In each block, subjects were presented with a new set of visual stimuli of set size *ns*, with set sizes between *ns=2* and *ns=5.* Each visual stimulus was presented 12-14 times in a pseudo-randomly interleaved manner (controlling for a uniform distribution of delay between two successive presentations of the same stimulus within [1:2\**ns*] trials, and the number of presentations of each stimulus), for a total of *ns*\*13 trials. At each trial, a stimulus was presented centrally on a black background (Figure 1). Subjects had up to 7 seconds to answer by pressing one of three buttons on the controller. Key press was followed by visual feedback presentation for 0.75 seconds, then a fixation period of 0.5 seconds before the onset of the next trial. For each image, the correct key press was the same for the entire block, and all feedback was truthful. Upon pressing the correct key subjects were shown “Correct” while pressing any other key led to “Try again!” Failure to answer within 7 seconds was indicated by a “No valid answer” message. Stimuli in a given block were all from a single category of familiar, easily identifiable images (e.g. colors, fruits, animals) and did not repeat across blocks. Participants were shown all stimuli at the beginning of each block, and encouraged to familiarize themselves with them prior to starting the learning block.

**Figure 1:**
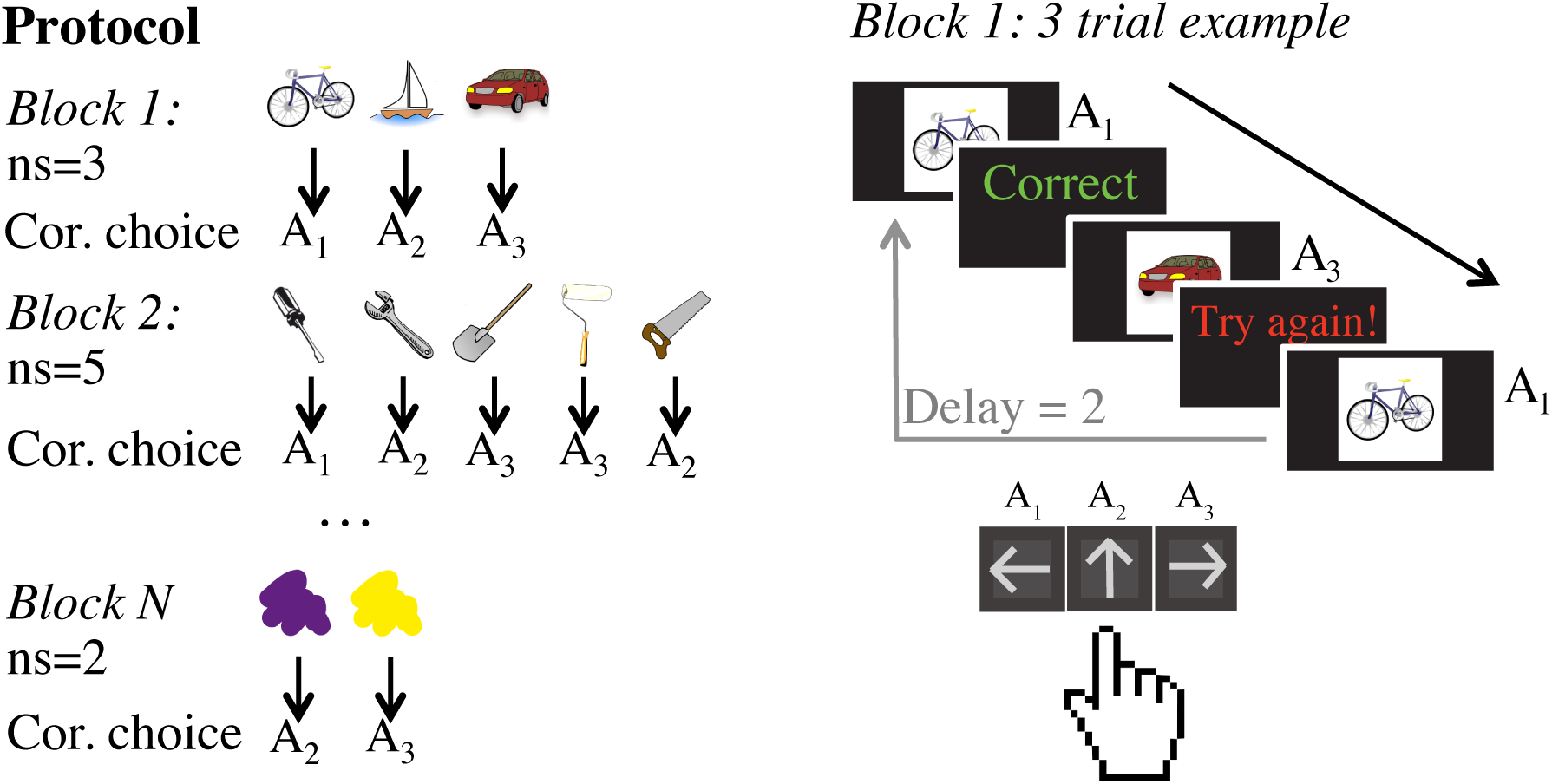
Experimental protocol. In each block, participants learned to select the correct action for each image for a new set of images. At the beginning of each block, the full set of images was presented for familiarization, then single trials began. On each trial of a block, participants responded to each stimulus by pressing one of three buttons on a hand-held controller. Immediately after responding they received deterministic truthful feedback (Correct/Try again!), before moving on to the next trial after a fixed interval. Crucially, participants needed to learn different numbers of stimuli (set size) in different blocks. Set size (ns) varied from 2 to 5.

We varied the set size across blocks: out of 10 blocks, 3 had set size *ns=2*, 3 set size *ns=3*, 2 set size *ns=4*, and 2 set size *ns=5*. We have shown in previous work that varying set size provides a way to investigate the contributions of capacity-and resource-limited working memory to reinforcement learning.

### Saliva collection

In addition to self-report measures of pubertal development, we also collected saliva from each of our subjects to quantify salivary testosterone. Testosterone is a reliable measure of pubertal status in boys and girls and is associated with changes in brain and cognition in adolescence ((Herting et al., 2014; Peper et al., 2011)).

Subjects refrained from eating, drinking, or chewing anything at least an hour before saliva collection. Subjects were asked to rinse their mouth out with water approximately 15 minutes into the session. At least one hour into the testing session, they were asked to provide 1.8 mL of saliva through a plastic straw into a 2 mL tube. Subjects were instructed to limit air bubbles in the sample by passively drooling into the tube, not spitting. Subjects were allotted 15 minutes to provide the sample. After the subjects provided 1.8 mL of saliva, or 15 minutes had passed, the sample was immediately stored in a −20°F freezer. The date and time was noted by the experimenter. The subjects then filled out a questionnaire of information which might affect the hormones measured in the sample (i.e. whether the subjects had recently exercised).

All subjects were asked whether they would like to complete two more saliva samples at home for additional compensation (another $25 Amazon gift card). Subjects who agreed to do the optional saliva samples were sent home with two 2 mL tubes, straws, and questionnaires identical to the one completed in lab. They were asked to complete each sample between 7:00 and 9:00 am on two different days following the testing session. Subjects were asked to refrain from eating, drinking, or brushing their teeth before doing the sample, and to fill out each questionnaire as soon as they were finished collecting saliva, taking care to note the date and time of the sample. Subjects were also instructed to keep the samples in the freezer, then wrap them in a provided ice pack in order to deliver the samples to the lab. Once both samples were complete, subjects contacted an experimenter and scheduled a time to return the samples, who gave the subjects their additional compensation, took note of any abnormalities in the samples, and immediately stored them in a −20 degree freezer. Samples in the −20 degree freezer were transferred weekly to a −80 degree freezer in an adjacent facility.

### Salivary Testosterone Testing

Salivary testosterone was quantified using the Salimetrics Salivatory Testosterone ELISA (cat. no. 1-2402, Bethesda, MA). Intra-and inter-assay variability for testosterone were 3.9% and 3.4%, respectively. Samples below the detectable range of the assay were assigned a value of 5 pg/mL, 1 pg below the lowest detectable value. Final testosterone sample concentration data were cleaned with a method developed by Shirtcliff and Byrne (in prep). Specifically, we produced a mean testosterone concentration from every salivary sample obtained from each of our subjects (every subject provided 1 to 3 samples). Subjects who had multiple samples below the detectable range of the assay (6 pg/mL) had their mean testosterone concentration replaced with 1 pg below the lowest detectable value (5). There were no subjects with any samples above the detectable range. Within subjects aged 8 to 17 only, outliers greater than 3 standard deviations above the group mean were fixed to that value, then incremented in values of +0.01 to retain the ordinality of the outliers.

### Model-independent analyses

We now describe how we analyzed the data from the behavioral task. To assess learning, we calculated the proportion of correct trials for each subject on each stimulus iteration. Each individual stimulus was repeated 13 to 14 times within a block. Within each set size, we calculated each subject’s average percentage of correct responses at each stimulus iteration (Figure 2a).

**Figure 2.**
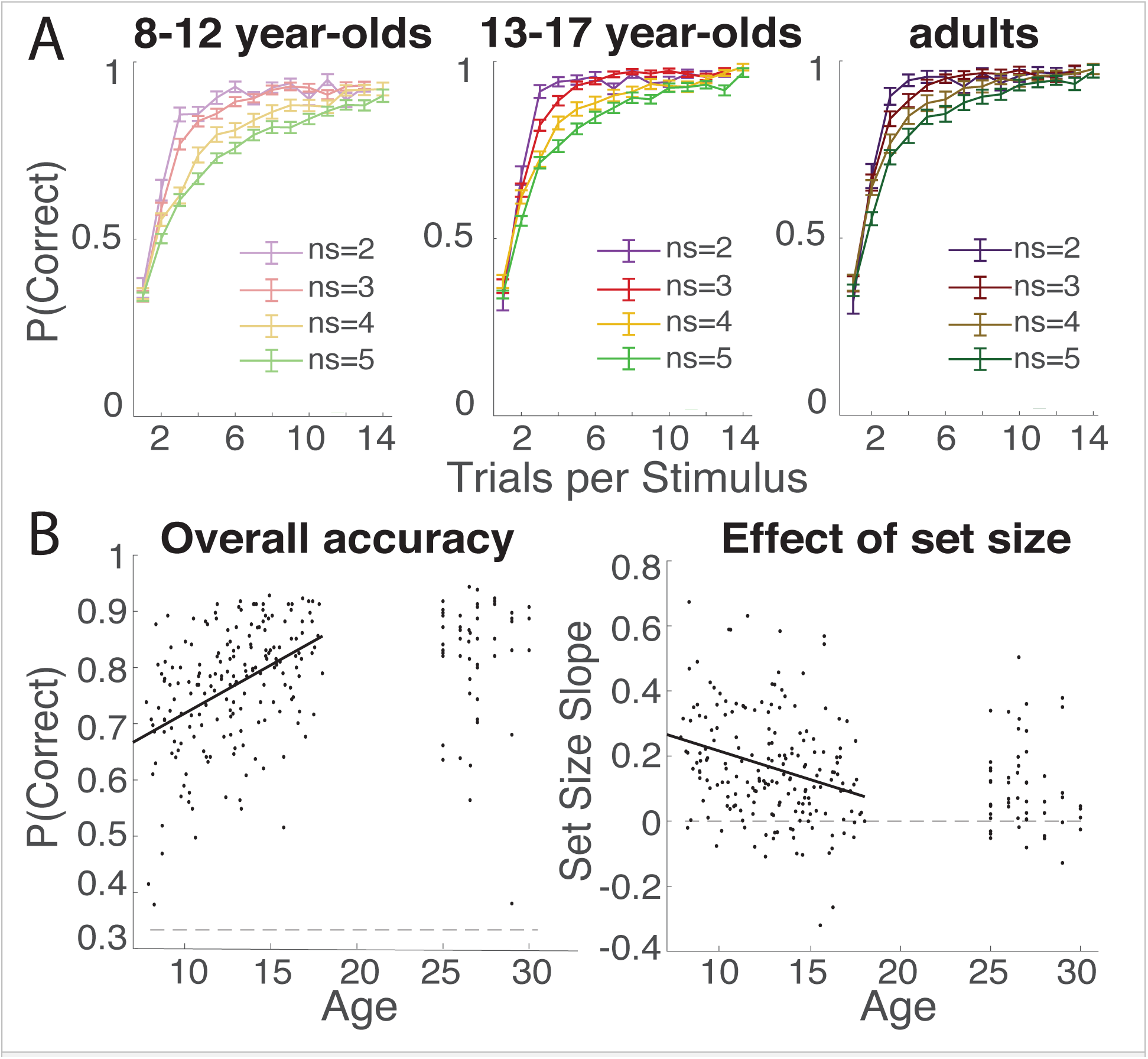
Age effects on behavior. **A.** Learning curves indicate the proportion of correct trials as a function of the number of encounters with given stimuli by set size (ns) in subjects aged 8 to 12, 13 to 17, and 25 to 30. All subjects quickly reached asymptotic accuracy in set sizes 2 and 3. In set sizes 4 and 5, learning was more graded. 8-12-year-olds appeared to learn slower than both 13-17-year-olds and adults (25-30). 13-17-year-olds and 25-30-year-olds exhibited similar learning. **B.** Overall accuracy and set size slope (linear effect of set size on performance; see methods) as a function of age. Each dot represents a single subject; lines indicate the best fit in a regression model. In subjects aged 8 to 17 overall accuracy significantly increased with age while set size effect significantly decreased, though it remained positive in all age groups.

We quantified the effect of set size on performance by calculating a set size slope (Figure 2b). The set size slope was a linear contrast of the form:

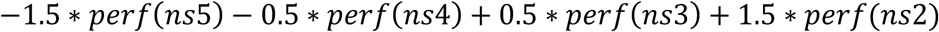

where perf(ns) is the average overall performance of trials within a block of set size ns.

Set size is correlated with delay between repetitions of the same stimulus, thus the previous measure may capture two separate aspects of the protocol that affect working memory (load and decay). In order to more precisely assess the effects of our set size manipulation on performance, we used a logistic regression to model trial-by-trial accuracy as a function of previous correct trials (pcor), previous incorrect trials (pinc), number of trials since the last presentation of the same stimulus (delay), and set size (ns) as predictors (Collins & Frank, 2012).

To understand the effects of development on behavior and on model parameters, we ran a series of analyses at group and individual levels. First, following the practice in the literature to separate “children” from “adolescents” and adults (Potter, Bryce, & Hartley, 2017) we grouped 8 to 12, 13 to 17, and 25 to 30-year-olds into separate groups and ran one-way ANOVAs on behavioral and modeling measures of interest, looking for broad group-based, age-related differences in cognition; ANOVAs were replaced by Kruskal-Wallis non-parametric tests for non-normal measurements. Where group effects were present, we ran post-hoc tests to identify which group drove the effect. We used t-tests when the measure was normally distributed, rank tests otherwise.

While helpful to visualize results, the binning into coarse 8 to 12 and 13 to 17-year-old groups for non-adult participants was arbitrary. We also analyzed the data of this 8 to 17-year-old subsample as a function of age as a continuous measure. Specifically, we ran non-parametric (Spearman) correlation tests on behavioral measures with age as a continuous predictor within the non-adult group.

Additionally, to further investigate the effects of age and pubertal development on our outcome measures, we examined relationships between behavior and model parameters as a function of pubertal development score (PDS) and salivary testosterone, either as continuous predictors, or defining binned groups (see supplementary material). Given that girls tend to begin puberty earlier than their male peers, and that the production of testosterone tracks the development of boys and girls differently, we also ran analyses of pubertal effects separating male and female subjects, and combining them. We also grouped participants aged 8 to 17 into narrower age bins based on quartiles. Further descriptions of how we grouped subjects into age, PDS, and testosterone bins, plus additional statistical test methods and results, are included in the supplemental material.

### Computational modeling

We used computational modeling to fit subject behavior and better quantify the separate involvements of reinforcement learning and working memory in task execution. We have shown that our model is able to separate the contributions of these two learning systems in both general ((Collins, 2018; Collins & Frank, 2012, 2018)) and specific populations ((Gold et al., 2017)). We tested 6 candidate models all built upon a simple reinforcement learning (RL) algorithm.

### Classic RL

The simplest model is a two parameter Q-learner (*RL*), which updates the learned value *Q* for the selected action *a* given the stimulus *s* upon observing each trial’s reward outcome *r*_*t*_ (1 for correct, 0 for incorrect):

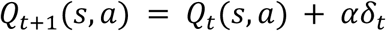

where *δ_t_* = *r*_*t*_ – *Q*_*t*_ (*s, a*) is the prediction error, and **α** is the learning rate, which is a free parameter. This model is similar to standard RL models as described in Sutton and Barto’s Reinforcement Learning: An Introduction ((2017)). Choices are generated probabilistically with greater likelihood of selecting actions that have higher *Q*-values. This choice is driven by a softmax choice policy, which defines the rule for choosing actions in response to a stimulus:

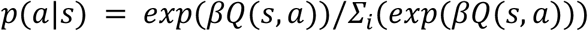

Here, **β** is the inverse temperature parameter determining the degree to which differences in Q-values are translated into more deterministic choice, and the sum is over the three possible actions a_i_.

### RL with undirected noise (RLe)

While the softmax allows for some stochasticity in choice, we also tested a model which allowed for “slips” of action. This was captured in an undirected noise parameter, *ε*. Given a model’s policy *π* = *p*(*a|s*), adding undirected noise consists in defining the new mixture choice policy:

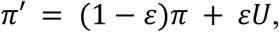

where U is the uniform random policy (U(a) = 1/n_A_, with number of actions n_A_ = 3). *ε* is a free parameter constrained to values between 0 and 1. This undirected noise captures a choice policy where with probability 1 – *ε* the agent chooses an action based on the softmax probability, and with probability *ε* lapses and chooses randomly. Failing to account for this irreducible noise can allow model fits to be unduly influenced by rare odd data points, like those that may arise from attentional lapses (Nassar & Frank, 2016).

### RL with perseveration (RLp)

To allow for potential neglect of negative feedback, we estimate a perseveration parameter *pers* such that for negative prediction errors (δ < 0), the learning rate *α* is reduced by *α* = (1 – *pers*)*α*. Thus, values of *pers* near 1 indicate perseveration with complete neglect of negative feedback, whereas values near 0 indicate equal learning from negative and positive feedback.

### RL with forgetting (RLf)

In this model we allow for potential forgetting of *Q*-values on each trial, implemented as a decay at each trial toward the initial, uninformed *Q-*value *Q*_*0*_:

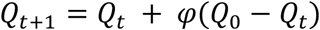

where 0 < *φ* < 1 is the forgetting parameter and *Q*_*0*_ = 1/n_A_.

### RL with 4 learning rates (RL4)

To improve the fit within the “RL only” class of models, we tested a version of the Q-learner that included a different learning rate *α* for each set size. Theoretically, this model could capture set size effects if they were driven by slower learning in higher set sizes.

### RL and working memory (RLWM)

This model incorporates two separate mechanisms by which learning can take place which interact at the level of choice. The first mechanism is a RL model as described above, with an inverse temperature parameter *β*, learning rate *α*, and undirected noise *ε*. The second mechanism is a working memory module.

The WM module stores weights between stimuli and actions, *W(s,a)*, which are initialized similarly to RL *Q*-values. To model fast storage of information, we assume perfect retention of the previous trial’s information, such that *W*_*t*+1_ (*s*_*t*_, *a*_*t*_) = *r*_*t*_, but include capacity limitation such that at most *K* stimuli can be remembered. To model delay-sensitive aspects of working memory (where active maintenance is increasingly likely to fail with intervening time and other stimuli), we assume that WM weights decay at each trial according to *W*_*t*+1_ = *W*_*t*_ + *φ*_*WM*_ (*W*_0_ – *W*_*t*_). The WM policy uses these weights in a softmax choice with added undirected noise, using the same noise parameters as the RL module.

Note that this implementation assumes that information stored for each stimulus in working memory pertains to action-outcome associations. Furthermore, this implementation is an approximation of working memory that assumes 1) rapid and accurate encoding of information when low amounts of information are to be stored; 2) decrease in the likelihood of storing or maintaining items when more information is presented or when distractors are presented; 3) decay due to forgetting. We believe these are key aspects of working memory.

RL and WM involvement in choice is modeled with a WM weight parameterized by the working memory capacity parameter K, and a WM confidence prior ρ. The overall choice policy is defined as a mixture using WM weight W_WM_ = ρ(min(1,K/ns)):

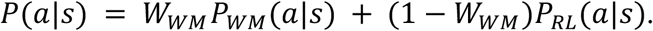

ρ captures the subject’s overall propensity to use WM vs. RL when within WM’s capacity. The WM weight then considers the capacity limit of the WM module as indicated by the proportion of items that can be maintained in working memory (min(1,K/ns)) as well as the subjects’ prior for relying on WM.

The strongest model in model comparison (by AIC score, figure 3b) was the RLWM model with six free parameters: the RL learning rate *α*, WM capacity *K*, WM decay *φ*, WM prior weight ρ, perseveration parameter *pers,* and undirected noise *ε*. The inverse temperature parameter *β* was fixed to 100, per previous experience showing that it allows best characterization of WM and RL parameters (Collins, 2018).

**Figure 3.**
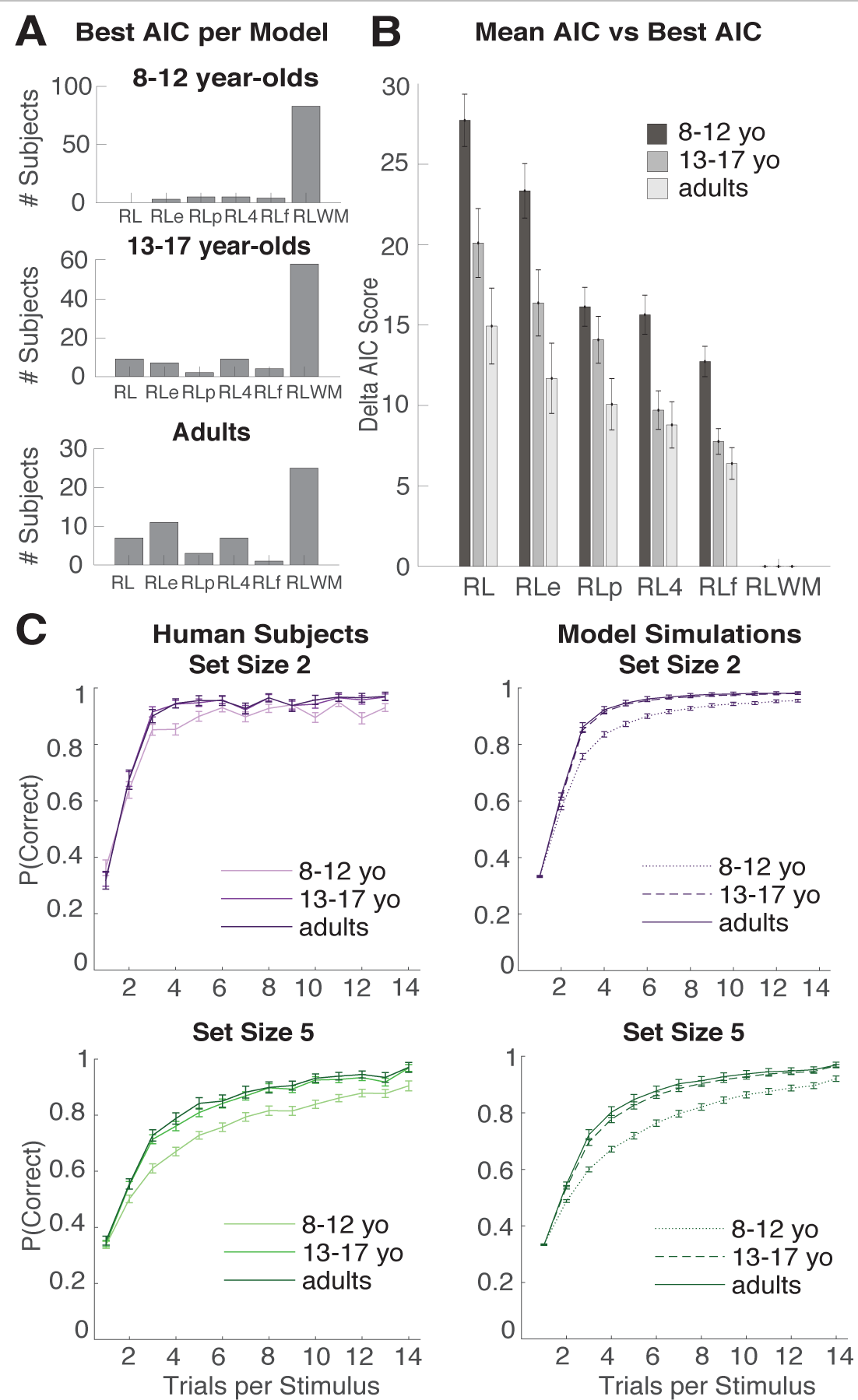
Model validation. **A.** Best fit model per subject. All subjects’ behavior was fit with 6 candidate models: reinforcement learning (RL), RL with an epsilon noise parameter (RLe), RL with perseveration (RLp), RL with four learning rates (RL4), RL with forgetting (RLf), and RL with working memory (RLWM; see methods). Plotted here is the number of subjects best fit by each candidate model in each age group as measured with AIC score. RLWM was the best-fitting model for a majority of subjects within subjects aged 8-12, 13-17, and 25-30. **B.** Difference in mean AIC score from the best fitting model (RLWM). Within subjects 8 to 12 years old (yo), 13 to 17 years old, and adults (25 to 30 yo), we calculated the mean AIC score for each candidate model, then compared to the mean AIC for the winning model (RLWM). Lower numbers indicate better fits. Error bars made with standard error of the mean. **C.** Model validation. Learning curves for participants (left) and model simulations (right) for set sizes 2 and 5 (see SI for all set sizes). RLWM model simulations with individual fit parameters accounted for behavior.

### Model fitting procedure

We used the Matlab constrained optimization function fmincon to fit parameters (the Mathworks Inc., Natick, Massachusetts, USA). This was iterated with 20 randomly chosen starting points, to increase the likelihood of finding a global rather than local optimum. All parameters were fit with constraints [0 1], except the capacity parameter *K*. Due to the non-continuous nature of *K,* each set of random starting points was paired with each of the possible fixed values [2 3 4 5] of *K*. The best fit within those possible values of *K* was selected as a proxy for optimizing *K* alongside the other parameters.

### Model comparison

We used the Akaike Information Criterion (AIC; Burnham & Anderson, 2002) to assess relative model fits and penalize model complexity. We previously showed that in the case of the RLWM model and its variants, AIC is a better approximation of model fit than Bayesian Information Criterion (BIC; Schwarz, 1978) at recovering the true model from generative simulations (Collins & Frank, 2012). Comparing RLWM and each of the variants of the simple RL model showed that RLWM provided a better fit to the data despite its additional complexity.

### Model simulation

Model comparison alone is insufficient to assess whether the best fitting model sufficiently captures the data, as it provides only a relative measure of model fit ((Wilson & Collins, submitted)). To test whether our models capture the key aspects of the behavior (i.e. learning curves), we simulated each model with fit parameters from each subject, with 100 repetitions per subject averaged to represent each subject’s contribution to group-level behavioral effects (Figure 3C).

## Results

### Overall accuracy

To analyze coarse age effects on behavioral and modeling measures, we first grouped participants into three groups by age (8 to 12-year-olds, 13 to 17-year-olds, and 25 to 30-year-olds), and tested the continuous effect of age on performance for non-adult participants.

All participants performed significantly better than chance (33%, Figure 2b). The mean accuracy for all subjects was 77.91% (median 80%). An ANOVA by age group revealed a main effect of group (F(240) = 20.29, p = 9.10e^-9^). Post-hoc t-tests show that this main effect was driven by the differences in performance of the 8-12 year-old group from the 13-17 year-old group (t(187) = 5.2, p < 10e^-4^) and the 25-30 year-old group (t(152) = 5.2, p < 10e^-4^). 8 to 12-year-olds’ performance was significantly worse overall (73% accuracy), while 13 to 17-year-olds and 25 to 30-year-olds performed similarly well (at 80.1% and 82.4% accuracy, respectively; t(141) = 1, p = 0.3). Within subjects aged 8 to 17, there was a positive correlation of age and overall accuracy (Figure 2b; Spearman ρ = 0.44, p = 2.e-10), confirming an expected improvement in learning performance with age.

### Learning

All age groups showed a similar qualitative pattern of learning whereby learning was faster in lower set sizes, a characteristic of combined working memory and reinforcement contributions to learning (Figure 2a). Learning reached asymptote for set sizes 2 and 3 within the first three or four trials, and reached asymptote incrementally in set sizes 4 and 5. There was a strong negative effect of set size on performance (set size slope; see methods; t(242) = 14.7, p = 2e-35; 207 of 243 participants). An ANOVA on the effect of set size revealed a main effect of age group (F(240) = 7.6, p = 0.0006). Post-hoc t-tests confirmed that this was driven by a larger negative effect of set size in 8 to 12-year-olds’ performance than in 13 to 17-year-olds’ (t(187) = 2.85, p = 0.0049) and adults’ (t(152) = 3.59, p = 0.0004). 13 to 17-year-olds were not more or less affected by set size than 25 to 30-year-olds (t(141) = 0.98, p = 0.33). Within subjects aged 8 to 17, there was a negative correlation between age and set size slope (ρ = −0.28, p = 8e-5; Figure 2b), supporting the previous analysis that participants’ learning became less sensitive to set size with increased age.

### Reaction time

Overall, mean reaction time (RT) decreased as a function of age (ρ = −0.34, p = 2e-6). RT variance also decreased as a function of age (ρ = −0.27, p = 0.0002). Reaction times were slower on high set size trials (t(242) = 33.1, p = 0; 242 out of 243 participants). There was no main effect of age group on the set size RT effect (F(240) = 1.24, p = 0.29).

All age groups and subjects appeared to be sensitive to the set size manipulation in both accuracy and reaction time, supporting the fact that all ages used both RL and WM to learn in this protocol. 8-to 12-year-olds were slower, less accurate, and more sensitive to the set-size manipulation than participants aged 13 and up. Performance rapidly reached adult levels by age 13 and remained stable until age 30. We next sought to use statistical and computational models to better characterize the underlying processes that drove the developmental changes in behavior.

### Logistic regression

For each subject, we ran a trial-by-trial logistic regression predicting response accuracy with predictors set size, delay since the current stimulus was last presented (two potential markers of WM), previous correct trials for that stimulus, and previous incorrect trials (two potential markers of RL; see methods). Set size had a negative effect on performance in all age groups (8 to 12-year-olds t = −6.96, p < 0.0003; 13 to 17-year-olds t = −4.39, p < 0.0003; 25 to 30-year-olds t = −3.88, p < 0.0003). Delay also had a negative effect on performance in all groups (8 to 12-year-olds t = −4.56, p < 0.0001; 13 to 17-year-olds t = −4.39, p < 0.0001; 25 to 30-year-olds t = −2.3, p = 0.026). Number of previous correct trials had a positive effect on performance in all age groups (8 to 12-year-olds t = 21.3, p < 0.0001; 13 to 17-year-olds t = 8.5, p < 0.0001; 25 to 30-year-olds t = 8.5, p < 0.0001). Number of previous incorrect trials had a negative effect in both children and teenagers but not in adults (8 to 12-year-olds t = −11.1, p < 0.0001; 13 to 17-year-olds t = −4.93, p < 0.0001; 25 to 30-year-olds t = −1.1, p = 0.27).

We tested the relationship between age and the individual logistic regression weights in a multiple regression predicting age from each individual’s five regression weights. After excluding subjects with weights 2 standard deviations above or below the mean (10 8 to 17-year-olds), we found that the weight of the fixed effect (t(173) = 7.1, p < 10e-4), and the weight of previous correct trials (t(173) = −2.9, p = 0.004) predicted age. None of the other predictors (pinc, delay, ns) predicted age (p’s > 0.09).

Results so far confirmed our prediction that younger participants would perform worse than older participants. There was evidence for successful RL recruitment in all subjects based on use of previous correct feedback, as well as evidence of WM recruitment based on set-size and delay effects. However, results so far were ambiguous as to whether

WM, RL, or both drove learning improvement with age. While a decrease in set size effect could hint at a WM effect, it is equally possible that worse performance in high set sizes in younger children could be due to worse RL (as hinted by the logistic regression results), which can support learning when WM is unavailable (e.g. at high set sizes). To clarify these findings, we next sought to model individual participants’ behavior with a mixture model capturing both RL and WM contributions to learning (see Methods).

### Modeling

Model comparison favored the RLWM model over other candidate models in all age groups (Figure 3ab; see methods). The exceedance probability in favor of the RLWM model was 1 in all groups (Rosa, Bestmann, Harrison, & Penny, 2010). RLWM was also the best model out of 6 candidate models for 83 out of 100 8 to 12-year-olds, 58 out of 59 13 to 17-year-olds, and 25 out of 54 25 to 30-year-olds (Figure 3a). Model simulations with fit parameters reproduced subject behavior, as well as differences between age groups (Figure 3c, supplementary figure S1).

We first investigated two noise parameters (Figure 4: Joint, epsilon and perseveration) and as expected, there was an effect of age group on the decision noise parameter epsilon (Kruskall-Wallis p = 0.025). Post-hoc comparison revealed that this was driven by the separation in behavior between children 8-12 and the older age groups (Rank-sum test 8 to 12-year-olds vs. 13 to 17-year-olds: p = 0.0003; 8 to 12-year-olds vs. 25 to 30-year-olds: p = 0.002; 13 to 17-year-olds vs. 25 to 30-year-olds: p = 0.94). There was a negative relationship between age and decision noise in the 8 to 17-year-old sample (ρ = −0.25, p = 0.0005), in boys (ρ = −0.22, p = 0.024), and in girls (ρ = −0.3, p = 0.005), showing that decisions were less noisy in older participants. There was also an effect of age group on the perseveration parameter (Kruskal-Wallis p = 0.0003). Post-hoc comparisons also revealed that the difference between 8 to 12-year-olds and the other age groups drove the main effect of group (Rank-sum test 8 to 12-year-olds vs. 13 to 17-year-olds: p < 10e-4; 8 to 12-year-olds vs. 25 to 30-year-olds: p = 0.001; 13 to 17-year-olds vs. 25 to 30-year-olds: p = 0.67). In the 8-17-year-old sample, there was also a negative relationship between age and perseveration (ρ = −0.3, p = 1.9e-07). This relationship was present separately in both boys (ρ = −0.3, p = 0.002) and girls (ρ = −0.44, p = 1.9e-05),

**Figure 4.**
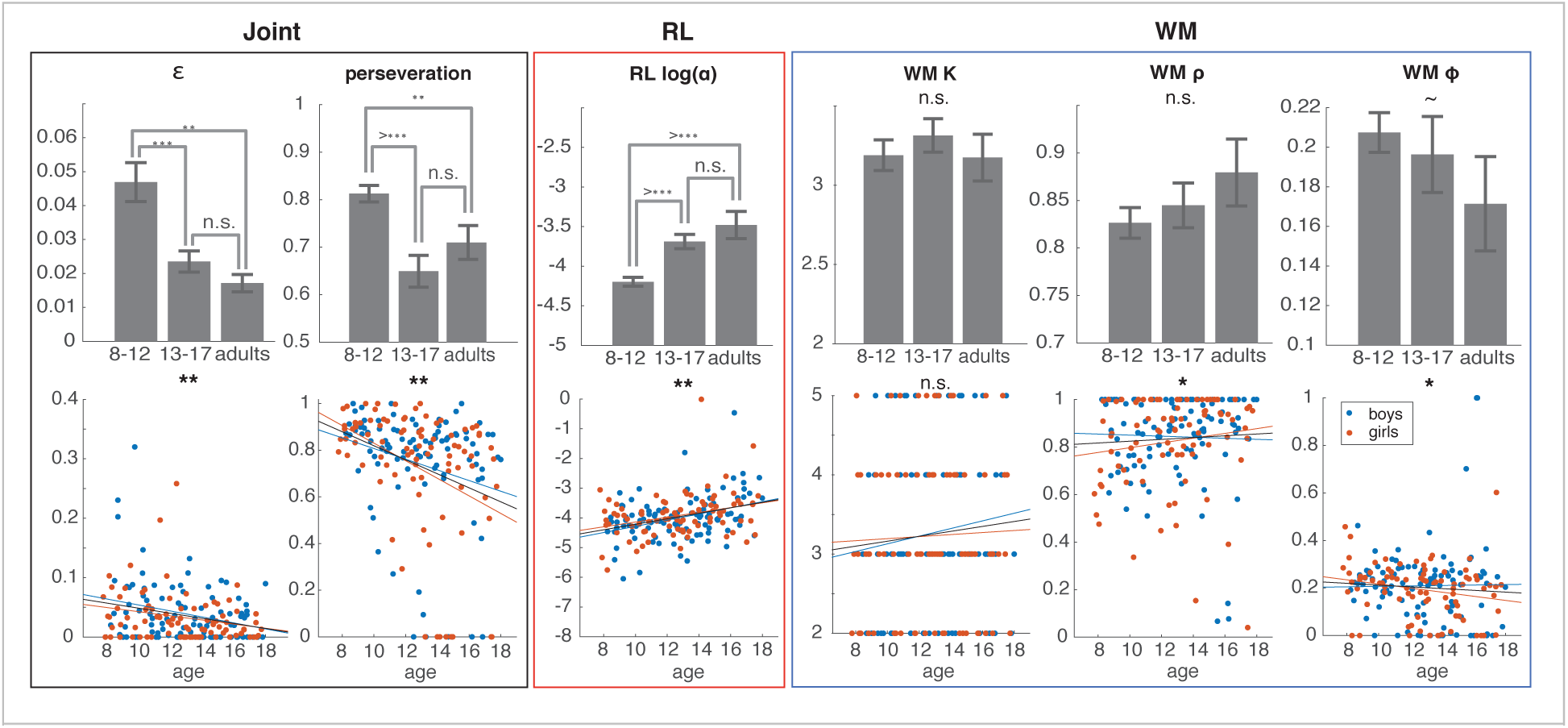
Effects of age on RLWM model parameters. There was an effect of both age group and age in years on the perseveration and ε decision noise parameters, whereby 8-12-year-olds had noisier behavior, and integrated negative feedback less than 13-17-year-olds and 25-30-year-olds. There were robust effects of age on the RL learning rate parameter α. There was no effect of coarse age groups or continuous effect of age within the 8-17 sample on the WM capacity parameter, and weak effects on WM weight ρ and decay ϕ. Error bars on bar graphs are standard error of the mean for each age group. ∼ indicates marginal significance at the p < 0.1 level, * indicates p < 0.05, ** indicates p < 0.01, n.s. stands for not significant.

We next investigated the relationship between age and RL learning rate α. There was a robust effect of age group on the RL learning rate parameter (Figure 4: RL; Kruskal-Wallis p = 0.0002). Additional comparison between groups showed that 8 to 12-year-olds had a lower average learning rate than 13 to 17-year-olds (p < 10e-4) and 25 to 30-year-olds (p < 10e-4). 13 to 17-year-olds did not differ from 25 to 30-year-olds (p = 0.3). Individual learning rates showed a significant upward trend with age 8-17 (ρ = 0.31, p = 2e-05), which was significant separately in both boys (ρ = 0.32, p = 0.001) and girls (ρ = 0.29, p = 0.007).

Finally, we investigated WM parameters. Surprisingly, we found no significant differences across our coarse age groups in WM capacity (Kruskal-Wallis test, p = 0.83), (Figure 4: WM left) (although see PDS and testosterone group results below). When we focused on individual subjects aged 8 to 17, we found no monotonic relationship between WM capacity and age (Figure 4: WM; Spearman ρ = 0.09, p = 0.23). We found no differences across coarse age groups in WM weight (Figure 4: WM; p = 0.25), but a marginal effect on WM decay (Figure 4; p = 0.08). These effects were stronger when investigating the continuous effect of age within non-adults: There was a small positive relationship between age and WM weight (Spearman ρ = 0.15, p = 0.04; Boys: ρ(101) = 0.06, p = 0.58; Girls: ρ(86) = 0.25, p = 0.02). There was also a negative relationship between age and WM decay (Spearman ρ = −0.17, p = 0.02) that was inconsistent across genders (Boys: ρ = −0.08, p = 0.42; Girls: ρ = −0.28, p = 0.01). This continuous effect was mostly driven by the youngest participants, with more WM decay and less WM weight.

Children of the same age can differ in their stage of pubertal maturation with considerable individual variability as well as sex differences in the timing of pubertal onset. Pubertal onset and later pubertal milestones may also produce non-monotonic changes over time such as a ‘U’ or ‘inverted U’ shape curve ((Piekarski, Boivin, et al., 2017; Piekarski, Johnson, et al., 2017)). Gonadal hormones, such as testosterone and its metabolites may also be playing an activational role in cortical or basal ganglia function on the day of testing ((Delevich, Thomas, et al., 2019)). Therefore, to explore developmental changes in learning with a finer resolution relevant to puberty we divided the 8-17 year old sample into 4 evenly divided bins first by age (<10.5, 10.6-12.8, 12.9-14.8, and >14.9 years old) then by pubertal development scale (PDS) (roughly pre-, early, mid-, and late/post-pubertal) and then by salivary testosterone (low, low-mid, mid-high, and high levels within gender) at time of test (T1) (see methods, Supplemental material). Cross-sectional data are quite limiting in differentiating age and pubertal effects, however we carefully explored the data and the effects of age, PDS, and testosterone concentration.

Division by finely graded age bins showed that noise and perseveration decreased with age (Supplementary Figures S4, S5; Noise: chi-squared = 15.69, p = 0.0013; pers: chi-squared = 24.1, p = 2e-05) and RL learning rate increased significantly with age (Supplementary Figure S6; chi-squared = 19.3, p = 0.0002) with no inverted U patterns. This general pattern was present in both boys (Supplementary Figures S4, S5, and S6; RL learning rate: chi-squared = 12.1, p = 0.007; noise: chi-squared = 6.7, p = 0.08; pers: chi-squared = 9.1, p = 0.028) and girls (RL learning rate: chi-squared = 8.1, p = 0.044; noise: chi-squared = 10.13, p = 0.018; pers: chi-squared = 26.2, p = 0.001). We observed similar results when non-adult participants were divided into bins based on PDS or testosterone measures (see supplementary table S2). Age, PDS and mean salivary testosterone were all also highly correlated (Supplementary Figure S2), therefore it was not clear if puberty onset played a significant role in RL related changes or not.

We next examined the relationship between finely graded age groups age, PDS and salivary testosterone measures and WM parameters. Here the greater number of groups enabled observation of ‘U’ or ‘inverted U shapes.’ Patterns of this type were apparent when subjects were grouped by PDS or T1. We had predicted potential effects of puberty onset on WM due to our previous work on effects of gonadal hormones on inhibitory neurotransmission in the PFC of rodent models.

(Piekarski, Boivin, et al., 2017; Piekarski, Johnson, et al., 2017). Subjects that were grouped by PDS did show significant differences in WM decay that was stronger in boys than girls (see Figure 5, Supplemental Figure 8; all: chi-squared = 10.02, p = 0.018; boys: chi-squared = 15.09, p = 0.0017; girls: chi-squared = 5.51, p = 0.14). Changes were strongest at puberty onset. We also uncovered a significant effect of testosterone at time of test (T1) on WM capacity (see Figure 5, Supplemental Figure 9; chi-squared = 13.15, p = 0.0043). This effect was marginal in boys alone (chi-squared = 6.61, p = 0.085) and significant in girls alone (chi-squared = 7.86, p = 0.049). When subjects were grouped by age (in 4 bins age 8-17), without regard to PDS or T, no significant group differences in WM parameters were found (though there was a marginal inverse U-shaped effect on WM capacity, and decreasing effect on decay; Figure 5, Supplemental Figures 8 and 9).

**Figure 5.**
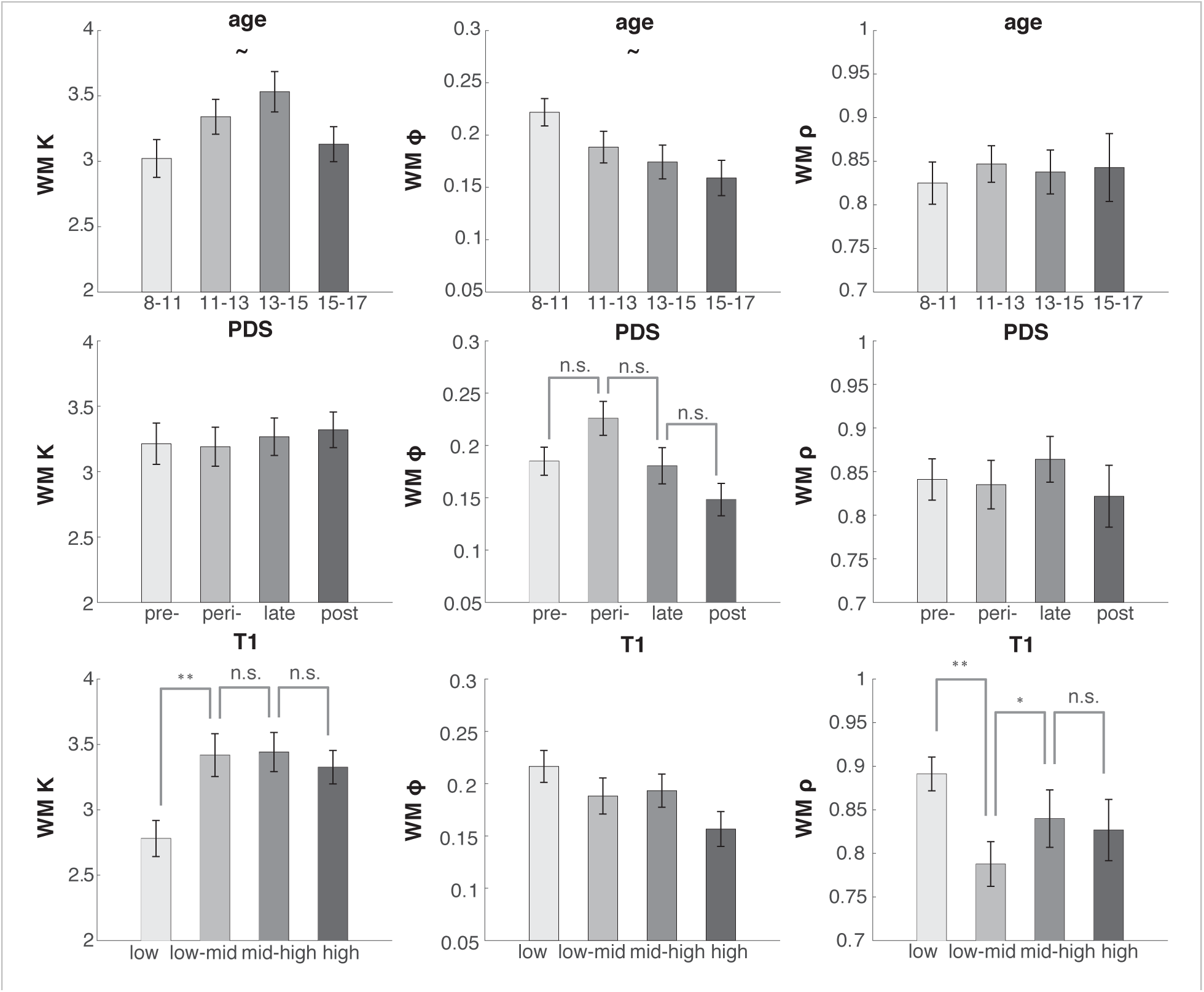
WM parameters by age, PDS, and sample 1 testosterone bins. All subjects aged 8 to 17 were binned according to age, pubertal development score (PDS), and salivary testosterone from the in-lab sample (T1). Girls and boys were binned separately according to gender-specific quartiles, then combined into equal-sized groups for each measure. Effects of group were assessed using the non-parametric Kruskal-Wallis test. Tests for which the group effect was significant at p < 0.05 were further examined with post-hoc non-parametric t-tests. n.s. indicates not significant, ∼ indicates marginal significance at p < 0.1 level, * p < 0.05, ** p < 0.01. Bin means are plotted; error bars show the standard error of the mean for each bin.

These results support our first prediction that changes in both RL and WM processes separately drive learning changes in subjects aged 8 to 30. However, we were surprised to find that changes in RL drive changes in learning more than changes in WM, in opposition to our second prediction. Finally, we support our third prediction that WM processes would differ in groups separated by pubertal development and testosterone.

## Discussion

In this study, we examined developmental changes in the working memory (WM) and reinforcement learning (RL) processes that contribute to simple stimulus-action association learning. While many developmental models emphasize late prefrontal cortex maturation and gains in WM in late adolescence ((Huizinga et al., 2006; Larsen & Luna, 2018)), previous studies have also shown changes in development in reinforcement learning tasks (Palminteri et al., 2016; Potter et al., 2017; Van Den Bos et al., 2012). However, these studies were not able to disentangle WM and RL contributions to learning. Our task and computational model were designed to address how WM and RL jointly contribute to learning in a simple task. ((Collins, 2018; Collins, Ciullo, Frank, & Badre, 2017; Collins & Frank, 2012; Collins et al., 2017)). The task and model are also notable because they reveal that WM is recruited in even simple reward learning tasks that are often assumed to include only model-free RL processes.

Here we used the RLWM task and computational methods to measure development of RL-and WM-based processes in children and teenagers aged 8 to 17 and adults aged 25 to 30. We found that 8 to 12-year-olds, 13 to 17-year-olds, and 25 to 30-year-olds all recruited both RL and WM in parallel for learning. 13 to 17-year-olds’ performance closely approximated 25 to 30-year-olds’, while participants aged 8 to 12 learned more slowly and reached a lower asymptotic accuracy. This was partially due to younger subjects’ noisier behavior, which may be indicative of a higher attentional lapse rate or tendency to explore different options, and to a lack of integration of negative feedback. However, we also related lower performance in younger subjects to a weaker contribution of RL to learning and to a smaller extent, to weaker WM maintenance and use.

Indeed, our behavioral results and computational model did not support our second hypothesis that WM changes are the main drivers of learning improvement from 8 to 17 years of age. Instead our data supported only our first hypothesis that RL-based learning develops from age 8 to 17, while some aspects of WM processes also did, though less strongly. Counter-intuitively, while 8 to 12-year-olds’ behavior was more sensitive to the learning load, we found that this was characteristic of a weaker RL system unable to make up for WM’s limitations in high set size conditions, more so than reflecting weaker capacity-limited WM. In addition to age effects on the decision noise and perseveration parameters, we found that the model’s RL learning rate (α) parameter increased with age; by contrast there were no significant effects of age on WM capacity. Decision noise and perseveration decreased with age while the RL learning rate steadily increased throughout adolescence and reached adult levels in 13 to 17-year-olds. Given the learning environment (a deterministic task with an unchanging reward structure), an optimal strategy would be to learn quickly from rewards (i.e. have a higher learning rate). This allows learners to reach asymptotic accuracy faster as long as the environment is stable. Thus 13 to 17-year-olds and 25 to 30-year-olds learned stimulus-action associations faster than 8 to 12-year-olds, which benefitted their overall task performance. We did observe small effects of age on WM weight and WM decay, reflecting increased use and stability of WM with age. Also, pubertal development and salivary testosterone concentations showed relationships with WM decay and capacity, respectively (at puberty onset).

We attempted to disentangle the influences of age and puberty on behavior despite the limitations of cross-sectional data and of our indirect measures of puberty, which is a complex multi-component process. Pubertal development was also highly correlated with age in our sample, however there were late and early developing individuals within our sample. When we examined noise, perseveration, and RL learning rate and divided subjects by pubertal development (PDS) or testosterone (T) concentrations, we largely observed the same patterns that were observed when using age. However, when we examined WM parameters in groups divided by PDS and T, we uncovered ‘inverted U’ shaped trajectories and found that WM decay changed significantly after pubertal onset in both boys and girls. WM capacity was also lowest in the group with the lowest testosterone concentrations at time of test. Thus pubertal status and gonadal hormone levels may also play a role in the development of WM decay and WM capacity, supporting our third hypothesis that there would be puberty-related changes in WM.

This study supports our first hypothesis but does not support our second hypothesis because we expected to observe strong age-related effects on WM processes. Instead, we observed the strongest age-related effects on RL learning rate. The strongest developmental effects on WM parameters were those of testosterone and pubertal development.

Adolescence is widely associated with the development of higher cognitive resources and the use of increasingly complex learning strategies ((Crone et al., 2006; Decker et al., 2016; Potter et al., 2017; Selmeczy et al., 2018)). These shifts in learning are mirrored by the development of prefrontal cortex, which does not reach functional or structural maturation until late in the second decade or even middle of the third decade of life ((Bunge & Gabrieli, 2002; Casey, Jones, & Hare, 2008)). Recent work proposes that adolescence as a life stage extends until age 25 ((Sawyer et al., 2018)). Furthermore, the use of WM in complex tasks has been shown to improve during development into late adolescence ((Crone et al., 2006; Geier et al., 2009; Huizinga et al., 2006; Luna, 2009; McAuley & White, 2011)). Therefore, it is somewhat surprising that we detected no age-related changes in WM capacity in subjects aged 8 to 17 despite our large sample. This is partially mitigated by PDS and T-related findings that show changes with the transition into puberty and the fact that other aspects of working memory (decay and weight) did show significant if weak effects of age. To better integrate this work into the established literature on the developmental science of learning, it may be helpful to dissociate the use of WM to maintain information from its use to manipulate information. The use of WM to manipulate information may develop over adolescence ((Crone et al., 2006)), while the use of WM to maintain information perhaps develops earlier. Recent results with a simple WM assay, a verbal span task, found age groups aged 9 to 25 performed equally well ((Potter et al., 2017)) and thus found no developmental effects on WM. The lack of age effect on the WM capacity in our study may reflect the fact that WM is used to maintain stimulus-action associations, rather than to manipulate information. However, in this task, information was never directly given to participants for them to hold – instead, they needed to integrate three temporally separate pieces of information (the stimulus observed, the choice made, and the feedback received), and determine the relevant information to maintain in WM. This, we would argue, constitutes an advanced mental manipulation in WM.

While dual systems models of development have often highlighted that subcortical areas mature before prefrontal cortex, there is also an important literature showing changes in striatal function through adolescence. Adolescents in the mid teen years have been shown more sensitive to and motivated by rewards than pre-pubertal children and post-pubertal adults ((Casey et al., 2008; Braams et al., 2015; Davidow, Foerde, Galván, & Shohamy, 2016; Gulley et al., 2018; Somerville, Jones, & Casey, 2010)). This increased motivation is reflected both in BOLD activation of the reward-responsive ventral striatum (Braams et al., 2015; A. Galvan et al., 2006b) and in increased recruitment of areas of the frontoparietal cognitive control network, when necessary ((Somerville & Casey, 2010;)). This could partially account for our results, explaining why 13 to 17-year-olds use RL more efficiently than 8 to 12-year-olds. However, such a relationship would predict differences between 13 to 17-year-olds and 25 to 30-year-olds, as seen in learning tasks that manipulate reinforcement more directly ((Palminteri et al., 2016)). We did not observe any difference between subjects aged 13 to 17 and adults in both behavior and in modeling; this might be due to a comparatively weak reinforcement manipulation, though it was sufficient to reveal strong differences between 8 to 12-year-olds and 13 to 17-year-olds.

An additional way to interpret our results might be to consider the role of episodic memory in our task. Recent research showed that a component of learning from reinforcement can be accounted for by episodic memory sampling ((Bornstein & Daw, 2012; Bornstein & Norman, 2017; Myers et al., 2002)). In fact, previous versions of this task found hippocampal contributions to learning ((Collins et al., 2017)). Thus, it is possible that some behavior captured by the RL component of the model actually included contributions of the episodic memory system, not just the reward learning systems. Some facets of episodic memory, like recognition memory, have been shown to develop by age 8, much earlier than prefrontal cortex-dependent executive function ((Ghetti & Bunge, 2012)). However, recent work shows that the interactions between hippocampus and prefrontal cortex – crucial for efficient memory search and retrieval – continue developing into adolescence ((Murty, Calabro, & Luna, 2016; Selmeczy et al., 2018)). Further, the development of both systems allows for the integration of relevant past experiences with goal-directed attention and can influence choice in the reward-driven BG ((Murty et al., 2016)). This developmental time course might account for some of the changes attributed to RL learning rate in our results. It will be an important topic for future research to better characterize episodic memory contributions to learning, in parallel to working memory and reinforcement learning’s contributions.

## Conclusions

Using a sample of 241 participants from 8 to 30 years old, we aimed to characterize the distinct developmental trajectories of multiple cognitive systems that contribute to learning in parallel, even in very simple situations: the reinforcement learning and working memory systems. Performance in a deterministic reward stimulus-action learning task which manipulated cognitive load revealed broad differences between subjects aged 8 to 12 and 13 to 17, whose performance was adult-like. While 8 to 12-year-olds were noisier in all conditions, their learning suffered more from a load increase than 13 to 17-year-olds’ and 25 to 30-year-olds’. In all participants, RL learning compensated for WM limitations under higher load. Computational modeling revealed that the stronger effect of load in 8 to 12-year-olds was less due to weaker time-limited, capacity-limited WM than to weaker RL compensation. This result was surprising based on the established literature on the late anatomical development of the prefrontal cortex. This work highlights the importance of carefully accounting for multiple systems’ contributions to learning when assessing group and individual differences and suggests that the development of reinforcement learning processes plays a major role in changes in learning during adolescent development. We hope these findings can fruitfully inform educational methods and intervention work. Future research in the science of learning should aim to develop experimental paradigms and computational models that more precisely dissociate the use of reinforcement learning, working memory, and episodic memory throughout development.

## Acknowledgements

We gratefully acknowledge the numerous people who contributed to this research: Lance Kriegsfeld, Celia Ford, Jennifer Pfeifer, Megan M. Johnson, Liyu Xia, Rachel Arsenault, Josephine Christon, Shoshana Edelman, Lucy Eletel, Haley Keglovits, Julie Liu, Justin Morillo, Nithya Rajakumar, Nick Spence, Tanya Smith, Benjamin Tang, Talia Welte, Lucy B. Whitmore, and Amy Zou. We are grateful to our participants and their families for their time and participation in this study.

## Funding

This work was funded by National Science Foundation SL-CN grant #1640885 to RD, AGEC, and LW.

The authors declare no conflict of interest.

## Supplementary Materials

### Supplemental Methods

#### Pubertal development scores

Subjects’ pubertal development scores (PDS) were assessed using the Petersen Development Scale ((Petersen & Crockett, 1988)). Subjects were asked to assess their pubertal development with 5 questions. Items assessed growth in height, growth of body hair, skin changes, and development of secondary sex characteristics. Subjects were asked to rate with one of four choices how development had progressed from “not started yet” to “has completed.” Ratings were assigned values of 1 −4. All answered questions were averaged to produce a PDS. The lowest possible PDS is a 1 while the highest possible is a 4.

#### Age, PDS, and Testosterone bins

To better understand the independent effects of age, pubertal development, and mean testosterone on model parameters, subjects aged 8 to 17 were binned into one of four groups for each measure. Age bins were created by splitting all subjects aged 8 to 17 into four quartiles (see Table 1). The age range (mean) of each bin is 8.06 to 10.54 (9.32), 10.62 to 12.85 (11.88), 12.92 to 14.86 (13.89), and 14.88 to 17.99 (16.31) years old.

**Table 1:** Subject N’s in each of four bins of age, PDS, mean testosterone, and testosterone from sample 1:

|  | low | mid | mid-high | high |
| --- | --- | --- | --- | --- |
| Age (M/F)<br>Mean (min-max) | 47 (26/21)<br>9.32 (8.06 – 10.54) | 47 (26/21)<br>11.88 (10.62 – 12.85) | 47 (26/21)<br>13.89 (12.92 – 14.86) | 46 (24/22)<br>16.31 (14.88 – 17.99) |
| PDS (M/F)<br>Mean | 47 (26/21)<br>1.19 | 42 (23/19)<br>1.86 | 45 (25/20)<br>2.80 | 47 (25/22)<br>3.59 |
| Mean T (M/F)<br>Mean (std) | 42 (23/19)<br>18.04 (5.78) | 44 (24/20)<br>33.65 (6.3) | 44 (24/20)<br>58.78 (17.19) | 47 (25/22)<br>102.80 (34.83) |
| T1 (M/F)<br>Mean (std) | 41 (22/19)<br>14.68 (4.79) | 43 (23/20)<br>26.04 (3.38) | 43 (23/20)<br>43.15 (9.19) | 46 (24/22)<br>100.49 (51.43) |

PDS bins were created in a similar fashion as the age bins. Subjects were split into gender-specific quartiles based on the range of PDS scores of all subjects of their gender, then combined into one bin. Since the PDS is derived from a questionnaire with 5 questions, there were many subjects with the same PDS. As such, the bins are not exactly equal in size (see Table 1). The mean PDS in each bin is 1.19, 1.86, 2.80, and 3.59.

Testosterone bins were also created with a quartile procedure. Within subjects under 18, subjects were split into gender-specific quartiles, one based on mean testosterone from 1 to 3 samples per subject and one based on testosterone concentration from sample 1, the in-lab sample taken in the middle of the experimental session. Subjects of the same gender-specific quartile (i.e. the males and females with the lowest levels of testosterone within their gender) were then binned together. The mean (std) testosterone levels of all mean testosterone bins were 18.04 (5.78), 33.65 (6.3), 58.78 (17.19), and 102.80 (34.83) picograms/mL. The mean (std) testosterone levels of the sample 1 testosterone bins were 14.68 (4.79), 26.04 (3.38), 43.15 (9.19), and 100.49 (51.43) picograms/mL.

### Supplemental Results

#### Pubertal development

To assess the specific effect of pubertal development on learning, we further tested the relationship between behavioral and modeling measures and pubertal development scale (PDS) and salivary testosterone. Age and PDS were highly correlated in adolescents aged 8 to 17 (Fig. S1, ρ = 0.92, p = 0), and in boys and girls separately (both ρ’s > 0.75, p’s < 1.e-19). Age and mean testosterone were also highly correlated in the whole sample (ρ = 0.52, p = 3e-17), in boys and men (ρ = 0.77, p = 2e-25), and in girls and women, though less so (ρ = 0.31, p = 9e-04). Testosterone and PDS were also highly correlated in all adolescents aged 8 to 17 and in both genders separately (Fig. S1, Adolescents: ρ = 0.47, p = 8e-22; Boys: ρ = 0.79, p = 3. e-21; Girls: ρ = 0.5, p = 2.1e-06). It is therefore difficult to disentangle effects of pubertal status from the effects of age in adolescents aged 8 to 18. Indeed, all analyses we conducted that included PDS or testosterone while controlling for age did not show effects of puberty. However, this could be due to the high correlation and does illuminate whether or not pubertal changes are potential causal variables. As such, we ran group and continuous analyses with each measure not accounting for the other measures and compared effects across measures.

The results of those analyses are presented in the main text. In summary, PDS results strongly support age results. There are also some PDS results in WM weight and decay. In contrast, testosterone results support age results for RL learning rate, noise and perseveration parameters, but relate to individual differences in WM capacity, in contrast to the effects on WM weight and decay for age and puberty measures.

#### Group Results

**Table 2:**
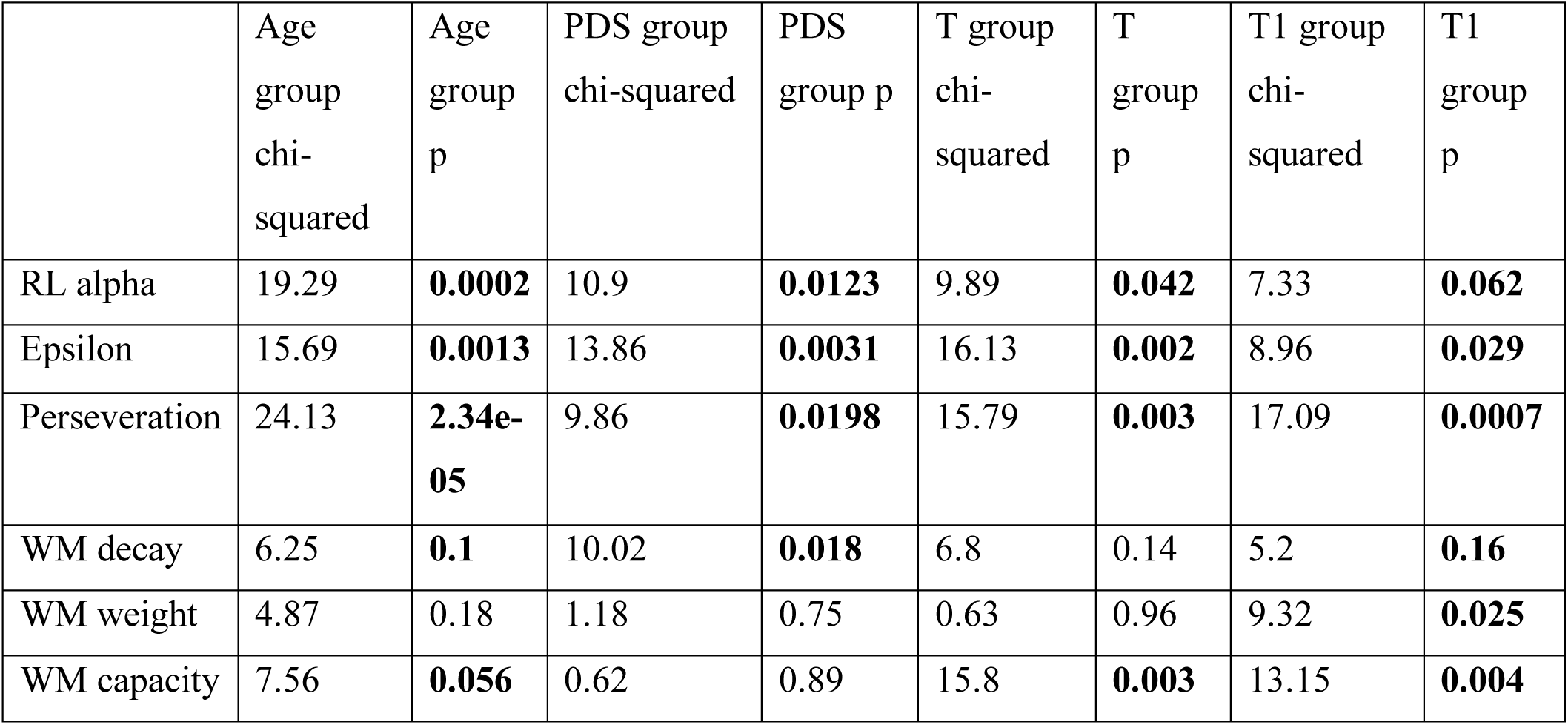
Kruskal-wallis tests on age groups, PDS, mean testosterone (T), and testosterone from sample 1 (T1) groups (all subjects aged 8 to 17):

|  | Age group chi-squared | Age group p | PDS group chi-squared | PDS group p | T group chi-squared | T group p | T1 group chi-squared | T1 group p |
| --- | --- | --- | --- | --- | --- | --- | --- | --- |
| RL alpha | 19.29 | **0.0002** | 10.9 | **0.0123** | 9.89 | **0.042** | 7.33 | **0.062** |
| Epsilon | 15.69 | **0.0013** | 13.86 | **0.0031** | 16.13 | **0.002** | 8.96 | **0.029** |
| Perseveration | 24.13 | **2.34e-05** | 9.86 | **0.0198** | 15.79 | **0.003** | 17.09 | **0.0007** |
| WM decay | 6.25 | **0.1** | 10.02 | **0.018** | 6.8 | 0.14 | 5.2 | **0.16** |
| WM weight | 4.87 | 0.18 | 1.18 | 0.75 | 0.63 | 0.96 | 9.32 | **0.025** |
| WM capacity | 7.56 | **0.056** | 0.62 | 0.89 | 15.8 | **0.003** | 13.15 | **0.004** |

A series of post-hoc non-parametric T-tests revealed that the lowest testosterone group exhibited smaller WM capacity than all other testosterone groups (p’s < 0.01). The low-mid, mid-high, and high testosterone groups exhibited no difference in WM capacity.

**Table 3:**
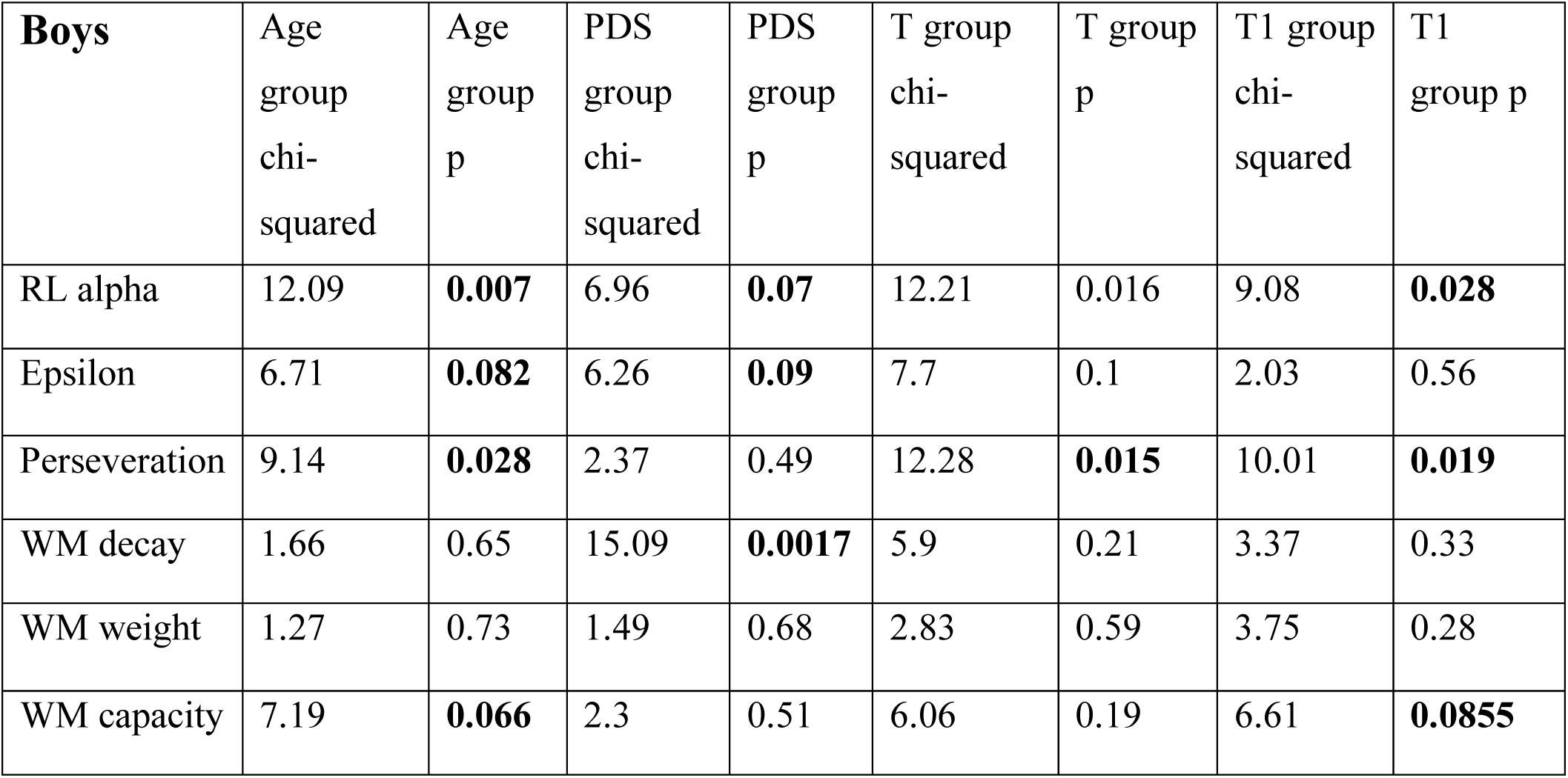
Kruskal-wallis tests on age, PDS, mean T, and T1 groups (male subjects only):

**Table 4:** Kruskal-wallis tests on age, PDS, mean T, and T1 groups (female subjects only):

| **Girls** | Age group chi-squared | Age group p | PDS group chi-squared | PDS group p | T group chi-squared | T group p | T1 group chi-squared | T1 group p |
| --- | --- | --- | --- | --- | --- | --- | --- | --- |
| RL alpha | 8.09 | **0.044** | 4.32 | 0.23 | 1.89 | 0.75 | 0.59 | 0.89 |
| Epsilon | 10.13 | **0.018** | 13.06 | **0.0045** | 11.61 | **0.021** | 10.24 | **0.017** |
| Perseveration | 26.17 | **0.001** | 10.44 | **0.0152** | 4.47 | 0.34 | 7.22 | **0.065** |
| WM decay | 5.6 | 0.13 | 5.51 | 0.13 | 4.57 | 0.33 | 5.37 | 0.14 |
| WM weight | 5.1 | 0.16 | 2.5 | 0.47 | 0.83 | 0.93 | 5.61 | 0.13 |
| WM capacity | 1.6 | 0.66 | 0.86 | 0.83 | 12.07 | 0.017 | 7.86 | **0.049** |

#### Continuous Results within 8-17 year-olds

**Table 5:** Spearman correlations of summary behavioral measures, and PDS, mean T, and T1:

| **Girls** | PDS rho | PDS p | Mean T rho | Mean T<br>p | T1 rho | T1 p |
| --- | --- | --- | --- | --- | --- | --- |
| Overall accuracy | 0.351 | **0.001** | 0.285 | **0.0025** | 0.343 | **0.0025** |
| Ns slope | -0.207 | **0.059** | -0.1901 | **0.045** | -0.148 | 0.12 |
| **Boys** |  |  |  |  |  |  |
| Overall accuracy | 0.324 | **0.0011** | 0.48 | **2.03e-08** | 0.464 | **1.39e-07** |
| Ns slope | -0.09 | 0.354 | -0.154 | 0.091 | -0.173 | **0.062** |

**Table 6:** Spearman correlations of model parameters, and age, PDS, and mean T in all subjects aged 8 to 17:

| Parameter | Age rho | Age p | PDS rho | PDS p | Mean T rho | Mean T<br>p |
| --- | --- | --- | --- | --- | --- | --- |
| RL alpha | 0.31 | **1.96e-05** | 0.21893 | **0.002** | 0.21 | **0.003** |
| Epsilon | -0.25 | **0.0005** | -0.269 | **0.0002** | -0.24243 | **0.001** |
| Perseveration | -0.37 | **2e-07** | -0.233 | **0.0015** | -0.26 | **0.0004** |
| WM decay | -0.16 | **0.019** | -0.11 | 0.12 | -0.09 | 0.21 |
| WM weight | 0.15 | **0.034** | 0.03 | 0.59 | 0.0103 | 0.89 |
| WM capacity | 0.08 | 0.23 | 0.05 | 0.49 | 0.17 | **0.021** |

**Table 7:** Spearman correlations of model parameters, and age, PDS, and mean T in girls and boys aged 8 to 17:

| **GIRLS (n = 87)** | **Age rho** | **Age p** | **PDS rho** | **PDS p** | **Mean T rho** | **Mean T p** |
| --- | --- | --- | --- | --- | --- | --- |
| RL alpha | 0.286 | **0.007** | 0.20 | **0.0677** | 0.11 | 0.3 |
| Epsilon | -0.3 | **0.005** | -0.33 | **0.002** | -0.332 | **0.002** |
| Perseveration | -0.44 | **1.963e-05** | -0.29 | **0.006** | -0.2031 | 0.066 |
| WM decay | -0.27 | **0.007** | -0.2005 | **0.069** | -0.159 | 0.15 |
| WM weight | 0.24 | **0.022** | 0.119 | 0.28 | -0.07 | 0.49 |
| WM capacity | 0.02 | 0.81 | 0.017 | 0.8745 | 0.11 | 0.3 |
| **BOYS (n = 100)** | **Age rho** | **Age p** | **PDS rho** | **PDS p** | **Mean T rho** | **Mean T p** |
| RL alpha | 0.319 | **0.001** | 0.211 | **0.036** | 0.32 | **0.001** |
| Epsilon | -0.223 | **0.024** | -0.228 | **0.023** | -0.19 | **0.061** |
| Perseveration | -0.304 | **0.002** | -0.112 | 0.27 | -0.32 | **0.001** |
| WM decay | -0.081 | 0.42 | 0.005 | 0.96 | -0.089 | 0.389 |
| WM weight | 0.06 | 0.48 | -0.04 | 0.67 | 0.071 | 0.48993 |
| WM capacity | 0.14 | 0.14 | 0.089 | 0.38 | 0.12 | 0.24 |

**Supplementary Figure 1.**
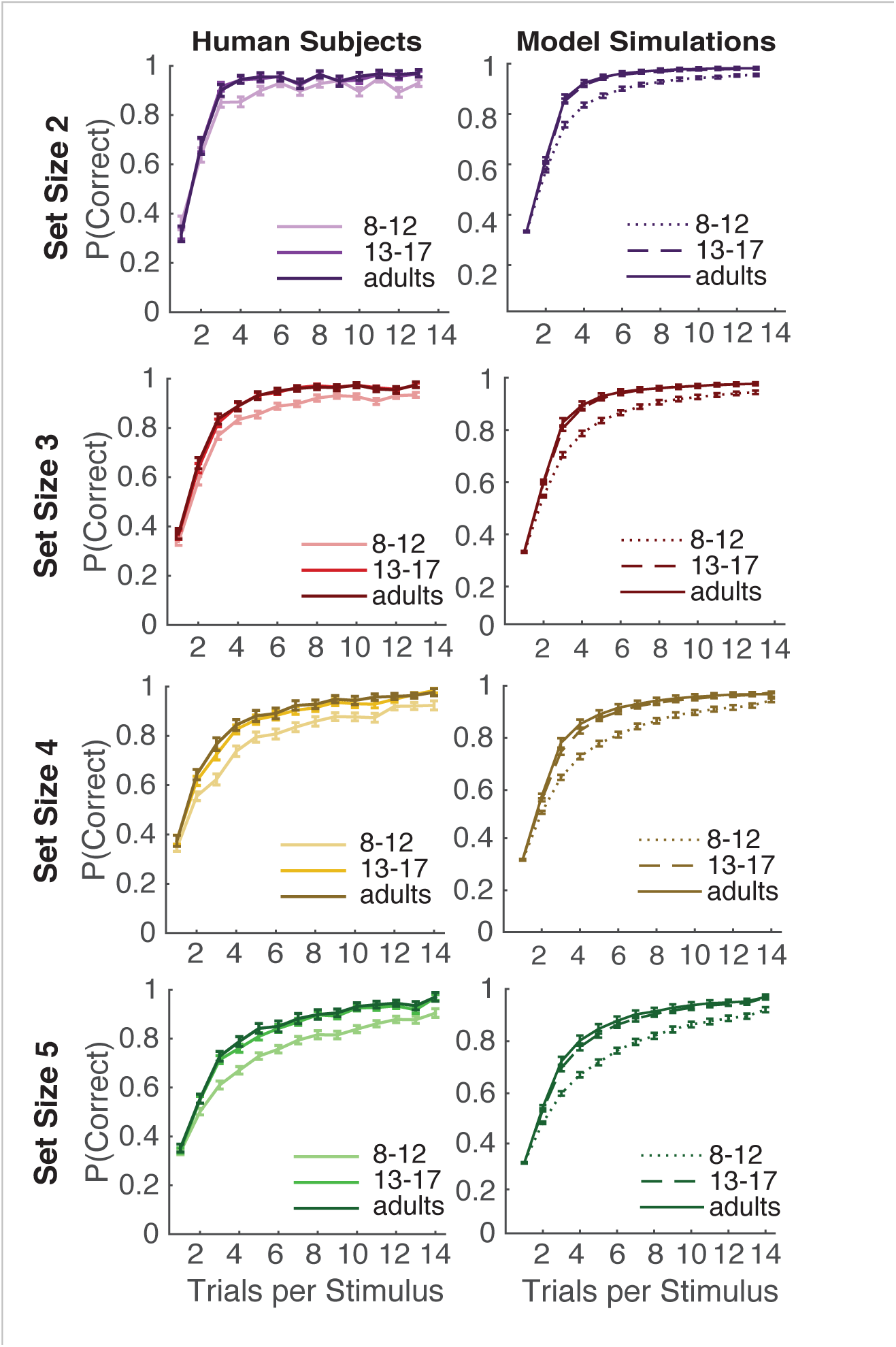
Model validation. Proportion correct per stimulus iteration in set sizes 2 – 5 for both human subjects and the RLWM model (see main Figure 3). Model simulation reproduces human behavior at all set sizes. The model can reproduce both age and set size effects when behavior is simulated with parameters fit to each subject individually.

**Supplementary Figure 2.**
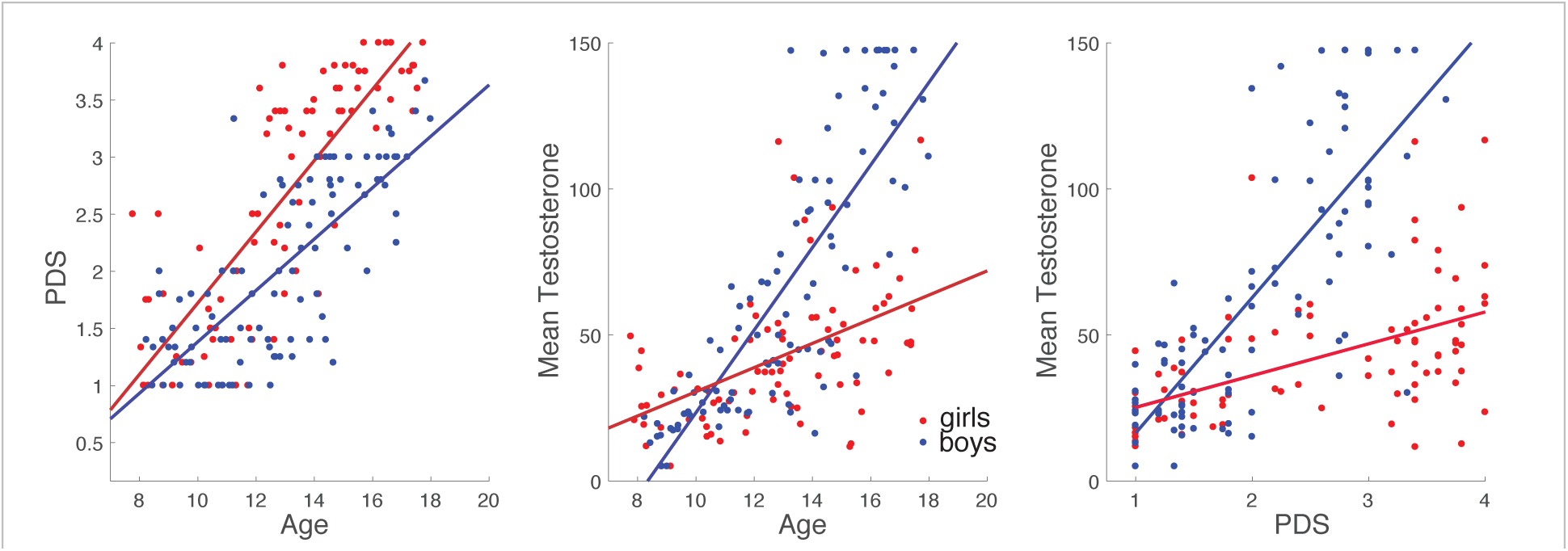
Relationships between puberty measures. In the full sample of adolescents aged 8 to 17, age, pubertal development score (PDS), and mean salivary testosterone are highly correlated. Each of these relationships also stands in boys and girls separately, though boys’ mean testosterone is more related to both age and PDS than girls’.

**Supplementary figure 3.**
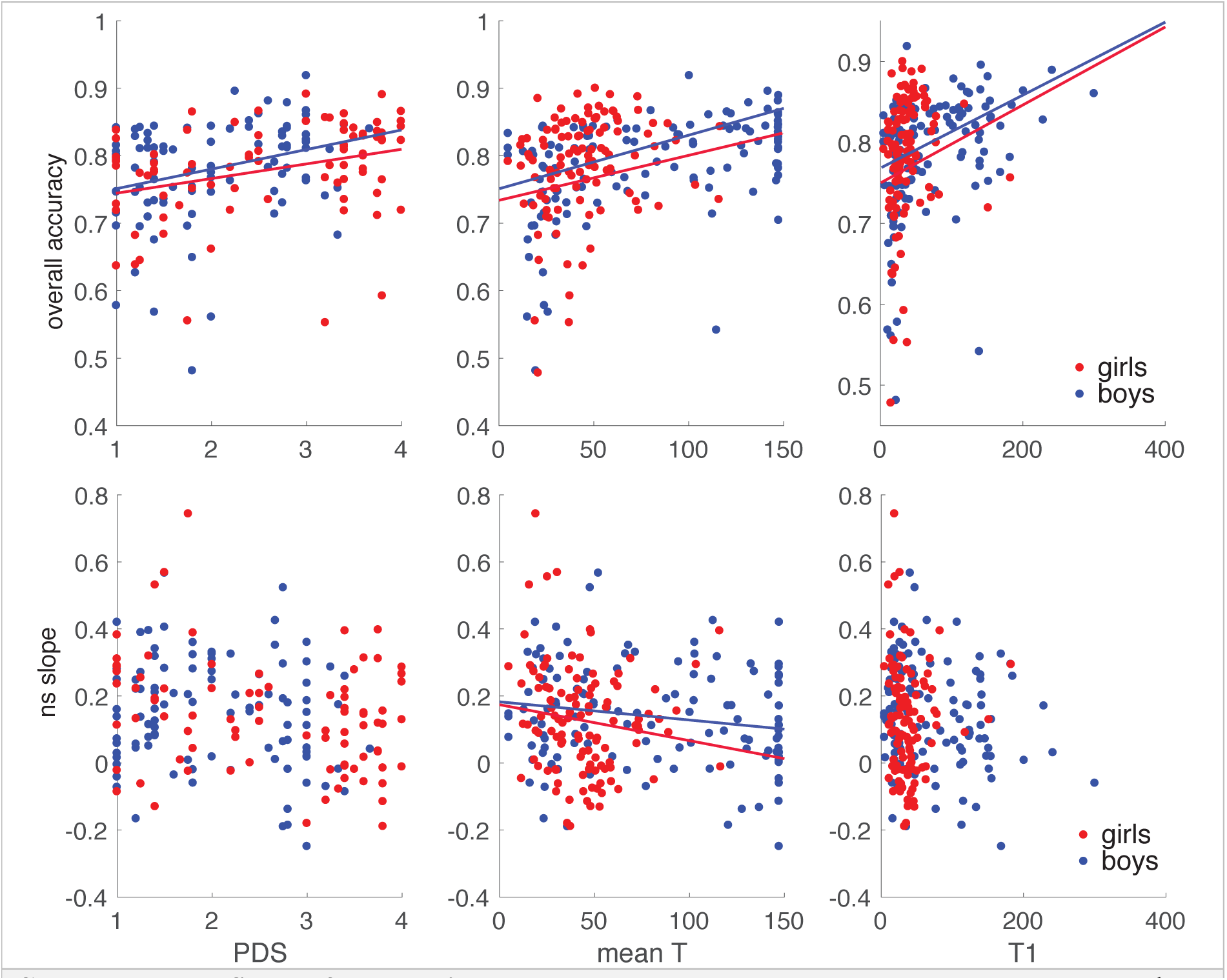
Behavioral summary measures by puberty measures. The relationships of pubertal development score (PDS), mean testosterone concentration across 1 to 3 saliva samples (mean T), and testosterone concentration in the in-lab sample (T1). Non-parametric Spearman correlations were run in girls and boys aged 8 to 17 separately. In girls, there was a positive effect on overall accuracy of PDS (ρ = 0.35, p = 0.0012), mean T (ρ = 0.29, p = 0.0025), and T1 (ρ = 0.34, p = 0.0025). The same was true in boys (PDS: (ρ = 0.32, p = 0.0011, Mean T: ρ = 0.48, p = 2.03e-08, T1: ρ = 0.46, p = 1.39e-07). The only effect of puberty on the set size effect (ns slope) in girls was that of mean T (ρ = −0.19, p = 0.046), though this effect was only marginal in boys (ρ = −0.15, p = 0.091). There was also a marginal relationship of PDS and ns slope in girls (ρ = −0.207, p = 0.059) and of T1 and ns slope in boys (ρ = −0.17, p = 0.062).

**Supplementary Figure 4.**
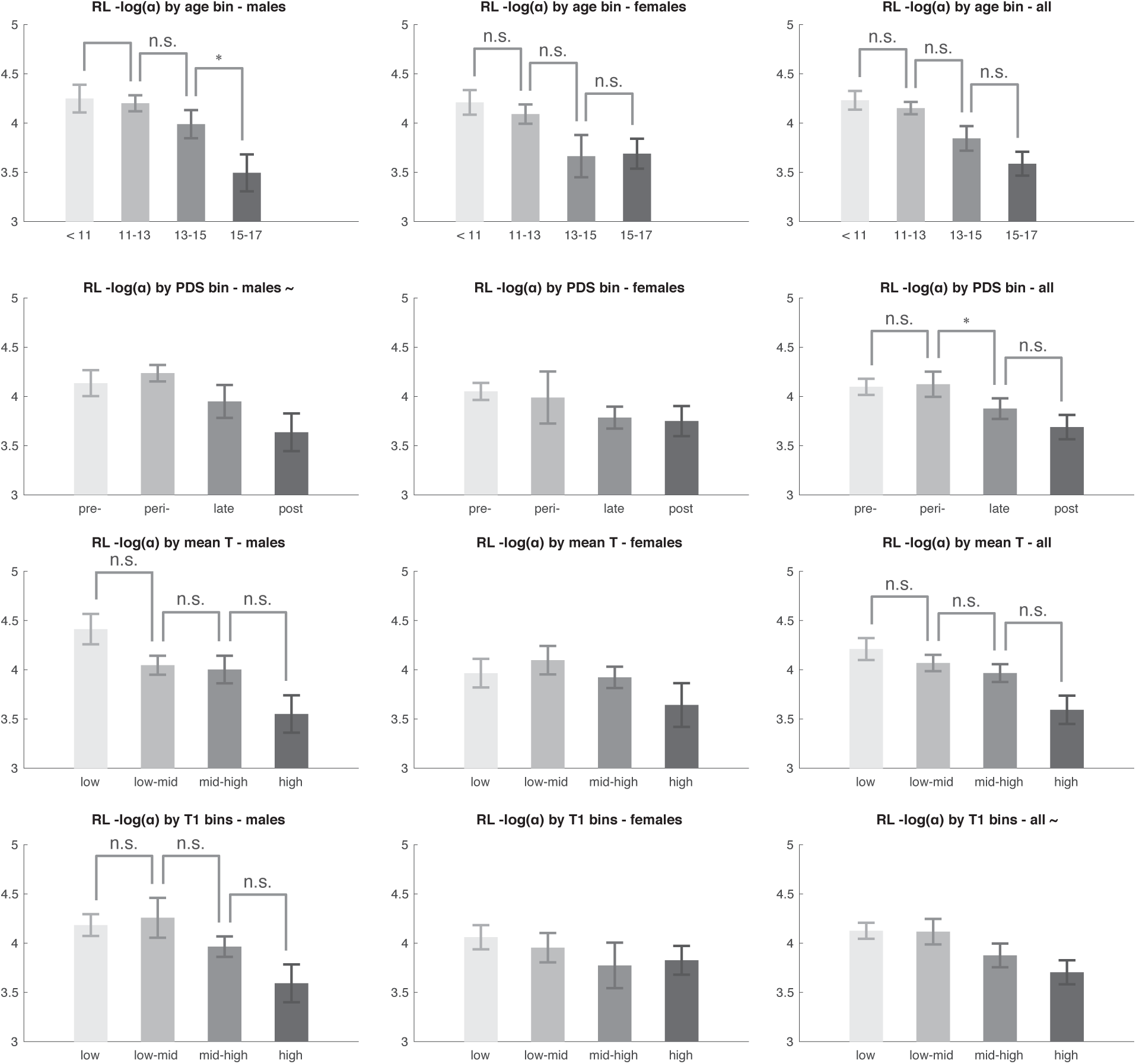
RL α (learning rate) by age, PDS, and testosterone (T) bins. Male and female subjects (8-17 years old) were binned into four groups using three different measures: age, PDS, or mean testosterone. Mean RL log(alpha) was then plotted for each of the four groups, in males alone, females alone, and the combined sample. For Kruskal-Wallis group p-values at or below a threshold of p = 0.05, post-hoc comparisons were run and their results plotted. For pairwise comparisons n.s. indicates not significant at p<0.05 threshold, * p < 0.05, ** p < 0.01 for pairwise comparisons; For Kruskal-Wallis tests ∼ indicates marginal significance at p < 0.1 level. Error bars represent the standard error of the mean. In both male and female subjects, we found significant group effects and PDS effects of age on learning rate.

**Supplementary Figure 5.**
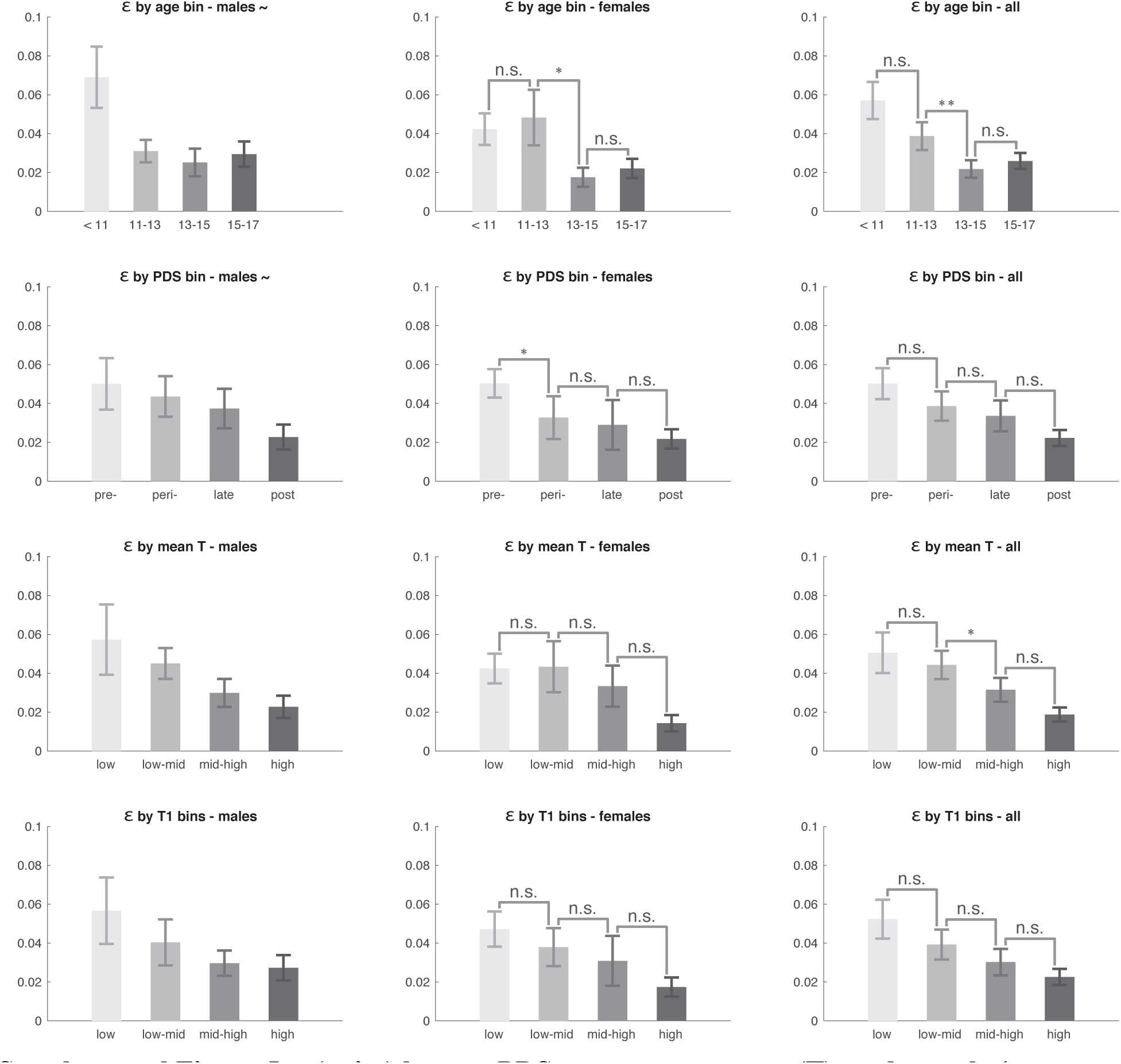
ε (noise) by age, PDS, mean testosterone (T), and sample 1 testosterone (T1) bins. Male and female subjects (8-17 years old) were binned into four groups using three different measures: age, PDS, or mean testosterone. Mean decision noise was then plotted for each of the four groups, in males alone, females alone, and the combined sample. For Kruskal-Wallis group p-values at or below a threshold of p = 0.05, post-hoc comparisons were run and their results plotted. For pairwise comparisons, n.s. indicates not significant at p<0.05 threshold, * p < 0.05, ** p < 0.01 for pairwise comparisons; For Kruskal-Wallis tests ∼ indicates marginal significance at p < 0.1 level. Error bars represent the standard error of the mean. We observed significant age, PDS, and T effects on epsilon in the entire sample, but only marginal effects in boys when separated from girls.

**Supplementary Figure 6.**
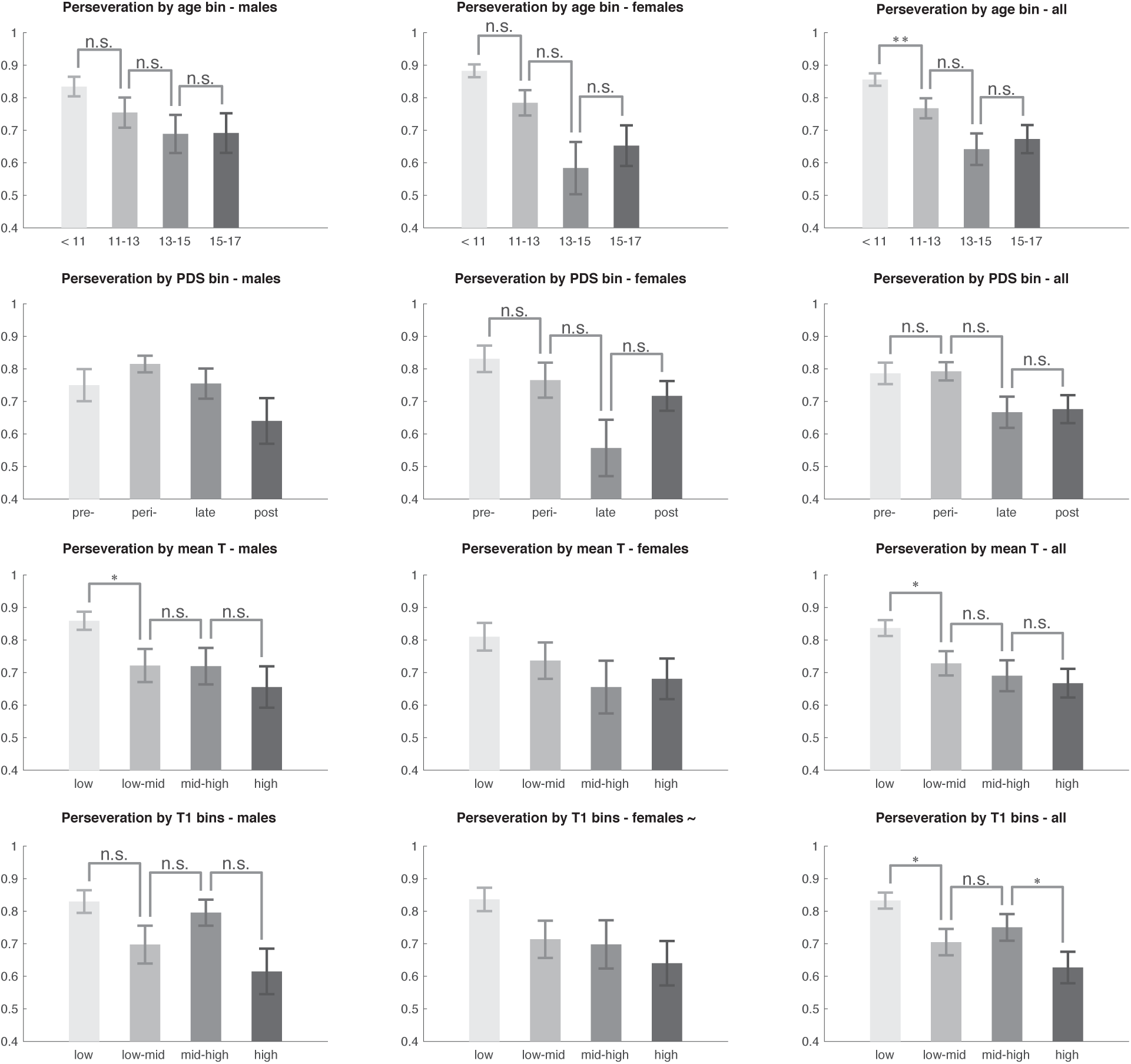
Perseveration by age, PDS, mean testosterone (T), and sample 1 testosterone (T1) bins. Male and female subjects (8-17 years old) were binned into four groups using three different measures: age, PDS, or mean testosterone. Mean perseveration parameter was then plotted for each of the four groups, in males alone, females alone, and the combined sample. For Kruskal-Wallis group p-values at or below a threshold of p = 0.05, post-hoc comparisons were run and their results plotted. For pairwise comparisons n.s. indicates not significant at p<0.05 threshold, * p < 0.05, ** p < 0.01 for pairwise comparisons; For Kruskal-Wallis tests ∼ indicates marginal significance at p < 0.1 level. Error bars represent the standard error of the mean. Perseveration decreased with age in a graded way across boys, girls, and the entire sample of 8 to 17-year-olds.

**Supplementary Figure 7.**
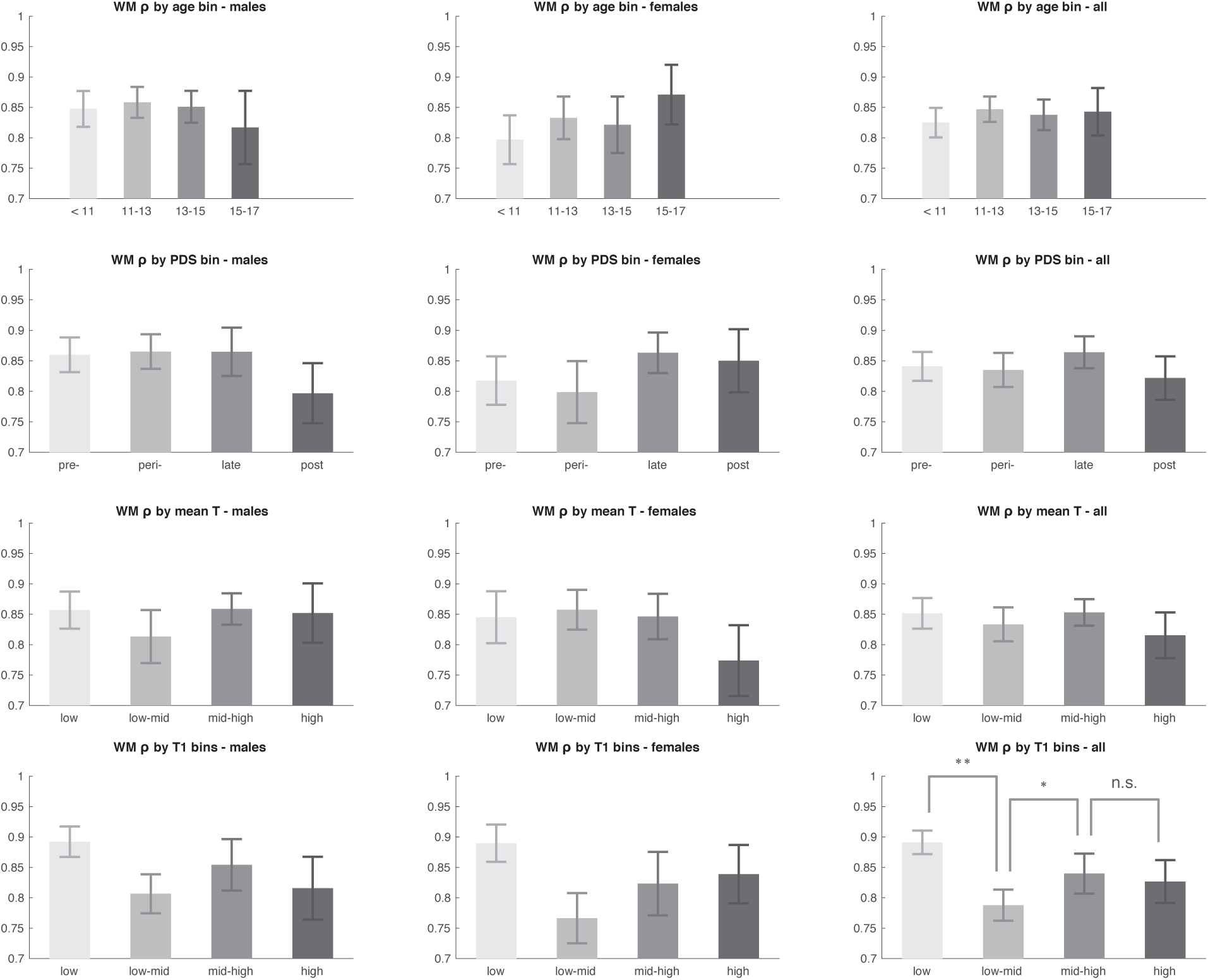
WM ρ (weight) by age, PDS, mean testosterone (T), and sample 1 testosterone (T1) bins. Male and female subjects (8-17 years old) were binned into four groups using three different measures: age, PDS, mean testosterone, or testosterone from sample 1 (T1). Mean WM weight was then plotted for each of the four groups, in males alone, females alone, and the combined sample. For Kruskal-Wallis group p-values at or below a threshold of p = 0.05, post-hoc comparisons were run and their results plotted. For pairwise comparisons n.s. indicates not significant at p<0.05 threshold, * p < 0.05, ** p < 0.01 for pairwise comparisons; For Kruskal-Wallis tests ∼ indicates marginal significance at p < 0.1 level. Error bars represent the standard error of the mean. We observed an effect of T1, activational testosterone, in the whole sample of 8 to 17-year-old subjects on WM use. There were no other development-related effects on WM use.

**Supplementary Figure 8.**
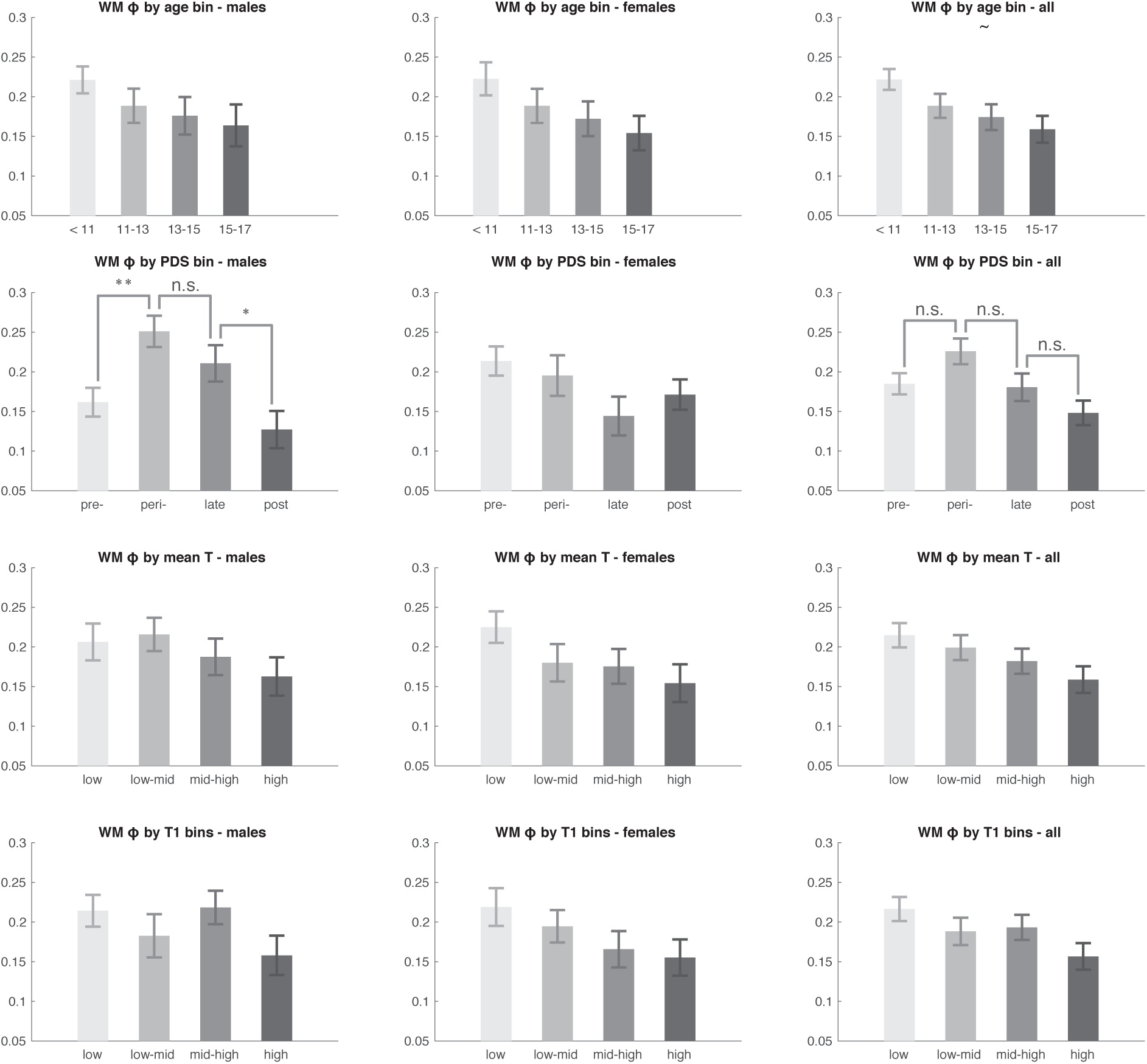
WM φ (decay) by age, PDS, mean testosterone (T), and sample 1 testosterone (T1) bins. Male and female subjects (8-17 years old) were binned into four groups using three different measures: age, PDS, mean testosterone, or testosterone from sample 1 (T1). Mean WM decay was then plotted for each of the four groups, in males alone, females alone, and the combined sample. For Kruskal-Wallis group p-values at or below a threshold of p = 0.05, post-hoc comparisons were run and their results plotted. For pairwise comparisons n.s. indicates not significant at p<0.05 threshold, * p < 0.05, ** p < 0.01 for pairwise comparisons; For Kruskal-Wallis tests ∼ indicates marginal significance at p < 0.1 level. Error bars represent the standard error of the mean. We observed an inverse U-shaped effect of pubertal development score (PDS) on WM decay in males and on the combined sample. Post-hoc PDS bin comparisons revealed that pre-and post-pubertal males appear to drive the overall effect of PDS on decay. The effect of age only reached marginal significance in the combined sample.

**Supplementary Figure 9.**
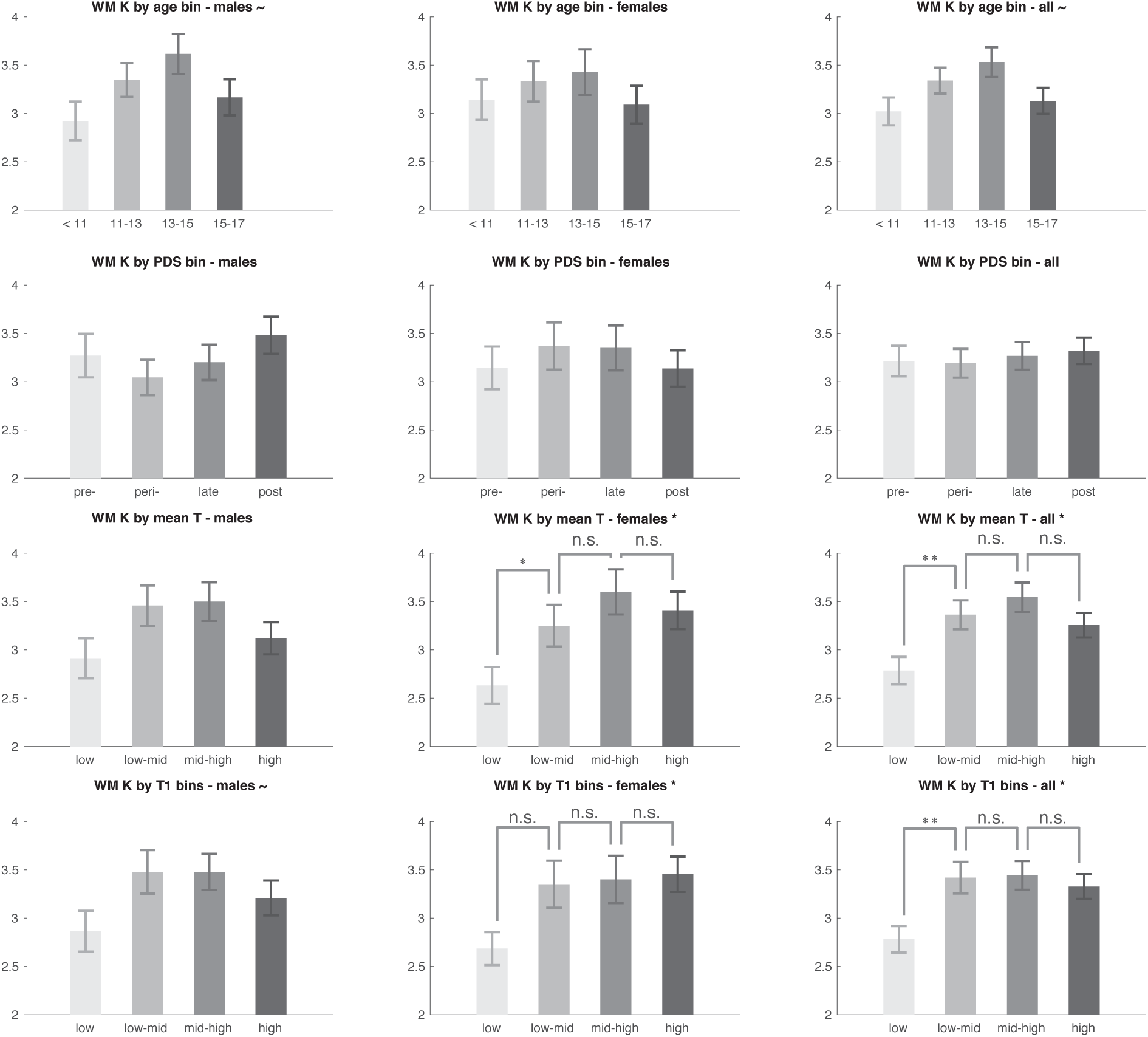
WM K (capacity) by age, PDS, mean testosterone (T), and sample 1 testosterone (T1) bins. Male and female subjects (8-17 years old) were binned into four groups using three different measures: age, PDS, mean testosterone, or testosterone from sample 1 (T1). Mean WM capacity was then plotted for each of the four groups, in males alone, females alone, and the combined sample. For Kruskal-Wallis group p-values at or below a threshold of p = 0.05, post-hoc comparisons were run and their results plotted. For pairwise comparisons n.s. indicates not significant at p<0.05 threshold, * p < 0.05, ** p < 0.01 for pairwise comparisons; For Kruskal-Wallis tests ∼ indicates marginal significance at p < 0.1 level. Error bars represent the standard error of the mean. There were robust effects of both mean testosterone and activational sample 1 testosterone on mean WM capacity. Closer examination of those effects revealed that subjects with the lowest level of sample 1 and mean T exhibited lower WM capacity.

